# Vitamin C functions as double-edge sword on cancer progression depending on ERK activation or inhibition mediated by its receptor SVCT2

**DOI:** 10.1101/2022.01.11.475954

**Authors:** Yian Guan, Bingxue Chen, Yongyan Wu, Zhuo Han, Hongyu Xu, Caixia Zhang, Weijie Hao, Wei Gao, Zekun Guo

**Affiliations:** College of Veterinary Medicine, Northwest A&F University, Yangling 712100, Shaanxi, China; Key Laboratory of Animal Biotechnology, Ministry of Agriculture, Northwest A&F University, Yangling 712100, Shaanxi, China; General Hospital, Clinical Medical Academy, Shenzhen University, Shenzhen 518055, Guangdong, China

**Author notes:** Correspondence. Prof. Wei Gao,. Prof. Zekun Guo,. These authors contributed equally (Yian Guan, Bingxue Chen, Yongyan Wu).

**Keywords:** Vitamin C, SVCT2, ERK, PTPN12, JAK2, GRB2, Oncotherapy

## Abstract

The effect of Vitamin C (Vc) in oncotherapy was controversial for decades. And hyperactivation of extracellular signal-regulated kinase (ERK) drove tumorigenesis. Herein, we demonstrated that Vc activated ERK through sodium-dependent Vc transporter 2 (SVCT2), while high-dose Vc resulted in persistent ERK feedback inhibition following activation. Extracellular Vc binding to SVCT2 initiated ERK activation, then transmembrane transport of Vc induced dimerization of SVCT2. Activated ERK phosphorylated protein tyrosine phosphatase non-receptor type 12 (PTPN12) at Ser^434^ and inhibited PTPN12 activity, thus enhancing phosphorylation of Janus kinase 2 (JAK2), which phosphorylated growth factor receptor bound protein 2 (GRB2) at Tyr^160^ to promote GRB2 dimers dissociation and recruitment of GRB2 to SVCT2, leading to further ERK activation. Different cancers have different sensitivities to Vc, the dose effects of Vc on cancer phenotypes depended on that ERK was activated or inhibited. These findings suggest SVCT2 is a Vc receptor mediating the ERK-PTPN12-JAK2-GRB2-ERK positive feedback loop and a potential target for oncotherapy.

**Abstract graphic:** 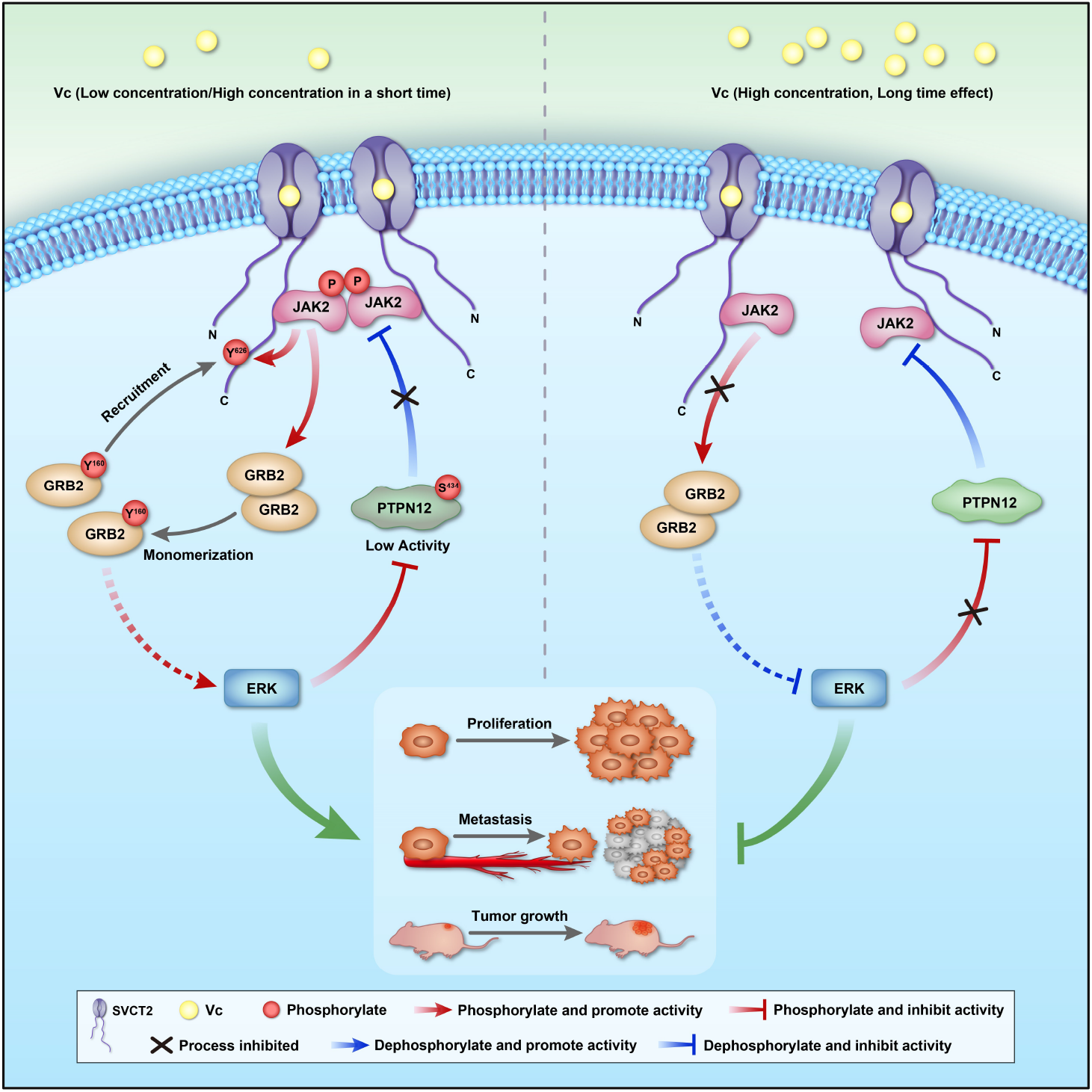

## Introduction

Vitamin C (Vc, also known as ascorbic acid or ascorbate) is an essential nutrient whose main physiological functions are to act as antioxidants and cofactors for various dioxygenases that participate in many physiological processes (Du et al., 2012). Meanwhile, Vc is also a controversial but widely used multifunctional drug molecule. Linus Pauling proposed in the 1970s that Vc has an anti-tumor effect. (Cameron and Campbell, 1974; Cameron and Pauling, 1974). Although subsequent studies showed no positive effects of orally administered Vc in cancer patients(Creagan et al., 1979; Moertel et al., 1985), numerous studies demonstrated that high-dose Vc at a millimolar range exerted anti-cancer effects(Padayatty et al., 2004; Verrax and Calderon, 2009). To date, two potential mechanisms have been used to describe the anti-cancer effects of high-dose Vc: (1) High-dose Vc can increase 5-hydroxymethylcytosine (5hmc) levels in cancer cells, altering the transcriptome and hence inhibiting malignant phenotypes(Mikkelsen et al., 2021; Peng et al., 2018). (2) High-dose Vc can promote H_2_O_2_ or ROS generation leading to DNA damage, thus killing cancer cells(Doskey et al., 2016; Su et al., 2019). However, the mechanism by which high-dose Vc inhibits cancer cells remains controversial(Ferrada et al., 2021), and it has been proposed that Vc may have unknown functions(Padayatty and Levine, 2016).

The mitogen-activated protein kinase (MAPK) pathway is one of the most conservative signal transduction pathways critical in cell proliferation, differentiation, migration, cell cycle transition, apoptosis and stress responses. Classical MAPKs mainly include ERK1/2, p38s, JNKs, and ERK5, while ERK3/4, ERK7/8, and NLK belong to atypical MAPKs(Sun et al., 2015). ERK1/2 is one of the most important MAPKs, and the RAS/RAF/MEK/ERK pathway is the most well-studied MAPK cascade. It has been reported that the hyperactivation of MAPK/ERK signaling exists in over 85% of cancer cases, and the RAS/RAF/MEK/ERK signaling is deregulated in over 40% of all types of cancer, which is mainly due to mutations in BRAF and its upstream activator RAS(Yuan et al., 2020). Increasing evidence indicates that the activation of ERK induces cancer cell proliferation, suggesting it is an important target for cancer therapy(Barbosa et al., 2021).

It has been reported that Vc activated the MAPK/ERK signaling pathway in acute myeloid leukemia cells and lung microvascular endothelial cells(Park et al., 2005; Varadharaj et al., 2006), and dehydroascorbic acid (DHA) was not involved in the regulation of MAPK/ERK signaling pathway activated by Vc(Ferrada et al., 2021). The mechanism of how Vc contributes to tumor treatment by affecting the MAPK/ERK signaling pathway is still unclear. Given its high prevalence in cancers, finding more upstream regulators and developing specific ERK inhibitors are the top priorities in screening anti-cancer drugs and treatment regimens.

Transceptor is a novel class of membrane proteins involved in environmental sensing, which possess dual activities in both acting as a transporter and initiating signal transduction like a receptor. So far, transceptors for amino acids, nucleosides, ammonium, phosphate, nitrate or sulfate have been identified in fungi, plants and mammals(Bröer and Gauthier-Coles, 2022; Maghiaoui et al., 2020; van den Berg et al., 2019; Van Zeebroeck et al., 2020; Zhang et al., 2020). Sodium-dependent Vc transporter 2 (SVCT2) was previously reported as a new member of transceptor that mediates JAK2 activation in sensing Vc(Han et al., 2021). Herein, we focused on the receptor activity of SVCT2 in mediating ERK activation induced by Vc, which was detected in a wide range of cell types. Upon binding to SVCT2, Vc induces ERK activation and following Vc transmembrane transport triggers dimerization of SVCT2, which is required for JAK2 trans-autophosphorylation. Furthermore, our study revealed that ERK, PTPN12, JAK2, and GRB2 form a positive feedback loop, which plays regulatory roles in the mutual interplay between ERK and JAK2 activation. Both low and high-dose Vc activated ERK, however, high-dose Vc resulted in persistent feedback inhibition following ERK activated violently. In addition, different cell types may have different sensitivities to Vc for subsequent ERK activation, the dose effects of Vc on cancer phenotypes depended on that ERK was activated or inhibited. These findings revealed a new Vc antitumor mechanism and highlighted the potential role of Vc in cancer prevention and control.

## Results

### Vc induces ERK activation in an SVCT2-dependent manner

Previously, we showed that Vc induced detectable JAK2 activation in F9 cells at 5 min after Vc supplementation and peaked at 30 min(Han et al., 2021). To assess the role of Vc in ERK signaling pathway, the F9 cells were treated with Vc for several periods. Strikingly, we found that phosphorylation of ERK was increased at 5 min after 50 μg/mL Vc treatment, peaked at 20 min, then transitioned to a decreased activation state at 30 min and persisted for at least 2 h, and finally disappeared after 12 h (Figure 1a and 1b). To further verify, transactivation assays were performed using pathway-profiling vector SRE-Luciferase (SRE-luc) as an indicator of ERK activity; the luciferase is driven by enhancer elements recognized by Elk-1/SRF, a downstream substrate of ERK(Buffet et al., 2015). Results confirmed that Vc increased the ERK signaling pathway activity (Figure 1c). Furthermore, Vc-induced ERK activation was also observed in HEK293T cells (Figure 1d and 1e), RAW264.7 cells (Figure S1a), and Jurkat T lymphocyte leukemia cells (Figure S1b). These results suggested that Vc-induced ERK activation is prevalent in multiple cell types.

**Figure 1.**
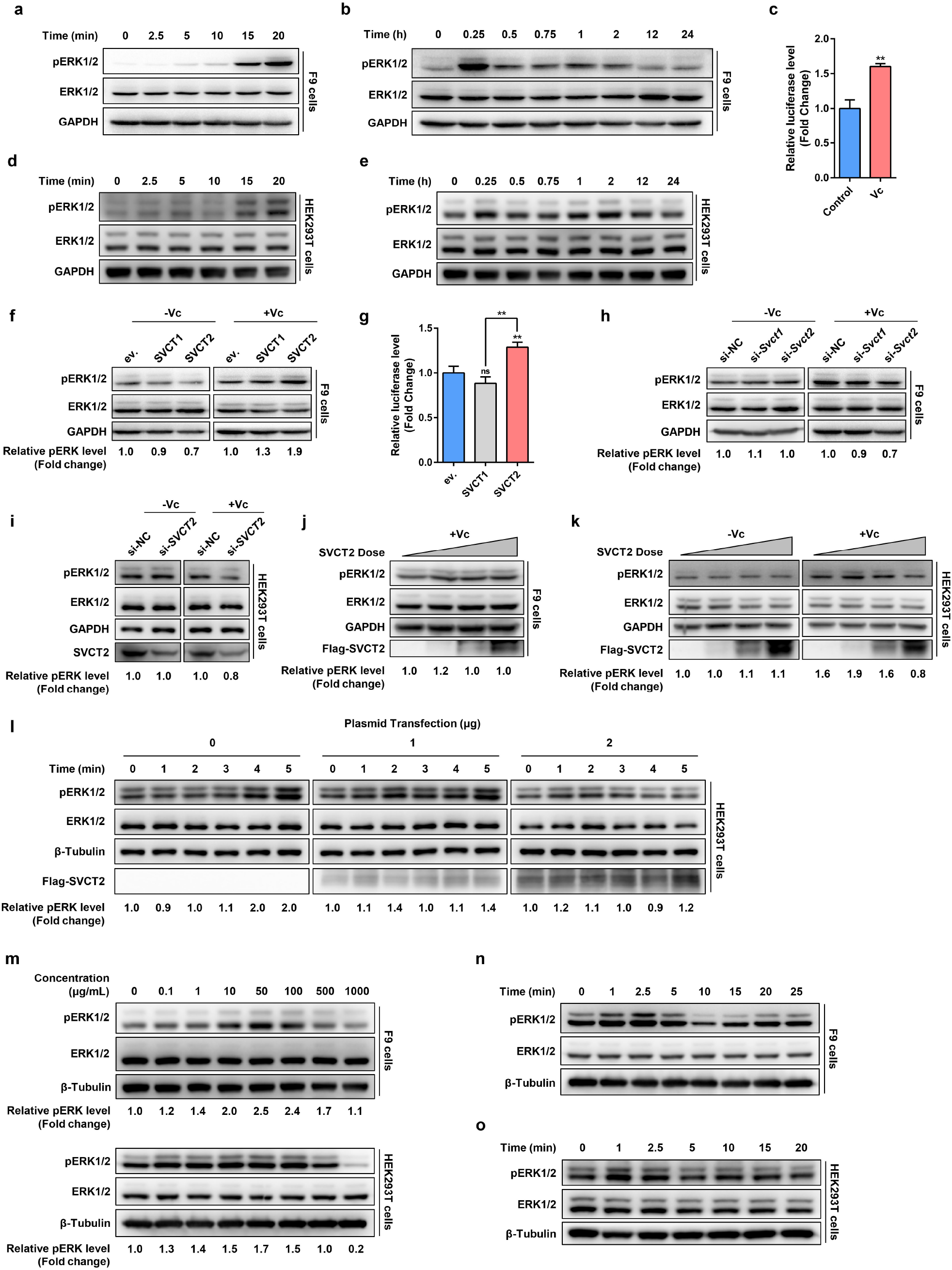
Vc induces ERK activation in an SVCT2-dependent manner. **(a and b)** F9 cells were treated with Vc (50 μg/mL) and lysed at indicated times. Endogenous pERK1/2, total ERK1/2 and GAPDH were detected by immunoblotting. **(c)** Dual-luciferase reporter (DLR) assay of F9 cells transfected with SRE-luc pathway-profiling vectors. 12 h after transfection, cells were incubated with Vc (50 μg/mL) and lysed 24 h later. Firefly luciferase activity was measured and normalized to *Renilla* luciferase. **(d and e)** HEK293T cells were treated with Vc (50 μg/mL) and lysed at indicated times. Endogenous pERK1/2, total ERK1/2 and GAPDH were detected by immunoblotting. **(f)** F9 cells were transfected with SVCT1 or SVCT2 plasmid as indicated for 36 h and then treated with Vc (50 μg/mL) for 20 min. Endogenous pERK1/2, total ERK1/2 and GAPDH were detected by immunoblotting. ev., empty vector. The expression of each vector was verified (data shown in Figure S1**c**). **(g)** DLR assay of F9 cells co-transfected with SRE-luc pathway-profiling vectors and indicated vectors. 12 h after transfection, cells were incubated with Vc (50 μg/mL) and lysed 24 h later. ev., empty vector. **(h and i)** F9 cells (**h**) or HEK293T (**i**) were transfected with *Svct1*-siRNA, *Svct2*-siRNA or negative control siRNA (si-NC) as indicated for 36 h and then treated with Vc (50 μg/mL) for 20 min, immunoblotting was performed as indicated. The knockdown efficiency was verified (data shown in Figure S1**d**-S1**f**). **(j and k)** F9 cells (**j**) and HEK293T cells (**k**) were transfected with different doses of Flag-SVCT2 plasmid for 36 h and then treated with Vc (50 μg/mL) for 20 min. Endogenous pERK1/2, total ERK1/2, β-Tubulin and Flag-tagged SVCT2 were detected by immunoblotting. **(l)** HEK293T cells were transfected with different doses of Flag-SVCT2 plasmid as indicated. 36 h after transfection, cells were treated with Vc (50 μg/mL) and lysed at the indicated times. Lysates were immunoblotted as indicated. **(m)** F9 and HEK293T cells were treated with Vc at indicated concentrations (0-1000 μg/mL) for 20 min and lysed immediately. Lysates were immunoblotted as indicated. **(n and o)** F9 (**n**) and HEK293T cells (**o**) were treated with Vc (500 μg/mL) and lysed at indicated times. Lysates were immunoblotted as indicated. Error bars represent SD of three independent experiments. **, P<0.01; ns, not significant; two-tailed Student’s *t*-test.

Our previous study also revealed that SVCT2 functions as a receptor-like transporter to mediate Vc-induced JAK2 activation(Han et al., 2021). To determine whether SVCTs mediated ERK activation induced by Vc, SVCT1 and SVCT2 were overexpressed in F9 cells, and phosphorylation of ERK was detected with and without Vc supplementation. Interestingly, we found that only exogenous SVCT2, but not SVCT1, could significantly enhance ERK activation in the presence of Vc, while the basal levels of pERK were not affected in the absence of Vc supplementation. (Figure 1f). Similarly, endogenous SVCT2 knockdown eliminated the effect of Vc on ERK activation (Figure 1h). In the Dual-Luciferase Reporter (DLR) assay, only SVCT2 overexpression increased ERK signaling pathway activity (Figure 1g). Similar results were also observed in HEK293T cells after SVCT2 knockdown (Figure 1i), supporting the possibility that SVCT2 mediated Vc-induced ERK activation.

To further assess the function of SVCT2 in mediating ERK activation, different doses of SVCT2 plasmids were transfected into F9 and HEK293T cells. Surprisingly, the phosphorylated ERK (pERK) levels were not enhanced at 20 min as the expression of SVCT2 increased after Vc stimulation and even exhibited gradual decrease in HEK293T cells (Figure 1j and 1k). Given that the activation of the signal pathway varied over time, the pERK level within 25 min induced by Vc was detected in SVCT2 transfected cells at different doses. Compared to the results in Figure 1d, two ERK activation cycles were observed due to the sampling time density, and the 20 min sampling point was actually in the second ERK activation cycle. In HEK293T cells transfected with an empty vector, the first activation cycle reached its peak at about 7.5 min, while that of cells transfected with 1.0 μg plasmid peaked at 5 min. However, the first ERK activation cycle induced by Vc was not observed in cells transfected with 2.0 μg plasmid (Figure S1g). Then, the first 5 min of Vc treatment were resampled with a sampling density of 1 min at an interval. As expected, in cells transfected with 2.0 μg plasmid, the first ERK activation cycle induced by Vc was maybe in 1 min or even earlier (Figure 1l), suggesting that the increased expression of SVCT2 in cells may shorten the ERK activation time in response to Vc. In addition, increased SVCT2 expression also resulted in a sharper wave of ERK activation in the first cycle, decreased level of pERK during interphase and a significantly weakened secondary activation cycle.

Next, pERK levels in F9 cells and HEK293T cells treated with various concentrations of Vc were detected. As shown in Figure 1m, the activation of ERK was observed in HEK293T and F9 cells treated with 0.1-1000 μg/mL Vc. Both showed the peak of ERK activation at Vc concentrations of 50 μg/mL and ERK activation effects was almost abolished in cells treated with 500 μg/mL Vc and was significantly inhibited in HEK293T cells treated with 1000 μg/mL of Vc. In order to reveal why high-dose Vc had no effects on ERK activation, the phosphorylation of ERK at various time scales was detected under 500 μg/mL of Vc supplementation. The results, consistent with SVCT2 overexpression results, showed that high-dose Vc still activated ERK; however, the activation time was earlier (2.5 min in F9 cells and 1 min in HEK293T cells), and followed by subsequent inhibition (Figure 1n and 1o). The long-term effect of high-dose Vc on ERK phosphorylation was also detected. In F9 cells, the pERK level was increased 0-2 h after high-dose Vc treatment, and began to decrease at 6 h, then showed significant inhibition at 12-24 h (Figure S1h). However, in HEK293T cells, the phosphorylation of ERK began to decrease at 1 h, and the significant inhibition lasted for at least 24 h (Figure S1i). Collectively, both low and high-dose Vc could activate ERK, and their differences may be mainly reflected in the activation time and the overall effect over a prolonged period. To exclude the possibility that pERK level resulted from elevated reactive oxygen species (ROS) induced by high-dose Vc, the ROS levels in HEK293T and F9 cells were detected by flow cytometry in real-time, and no significant change of the ROS levels was observed in the first 20 min after Vc supplementation (Figure S1j).

ERK signaling pathway plays an important role in the initiation and progress of cancer(Roberts and Der, 2007). Since the phosphorylation of ERK is affected by SVCT2 and Vc both in a dose-dependent manner, we wondered whether SVCT2 expression levels and Vc dose were relevant to cancers. Kaplan-Meier analysis showed that LIHC, KIRC, PAAD, LGG and BLCA patients with high SVCT2 expression level had longer overall survival, meanwhile high Vc levels were found in their corresponding tumor tissues (Figure 2a)(Padayatty and Levine, 2016). However, the opposite result was observed in ESCA patients with unavailable Vc concentration in esophageal tissue. Moreover, there was no correlation between SVCT1 expression level and overall survival in above cancers (Figure S2). Then the differential expression of SVCT2 (SLC23A2) in cancers and Vc concentrations in their corresponding tumor tissues was analyzed. The results showed that the expression of SVCT2 was abnormally expressed in cancers listed in Figure 2b. Intriguingly, in tissues with high Vc concentrations, such as adrenals and uterus(Basu et al., 1990; Luck et al., 1995; Padayatty and Levine, 2016), SVCT2 was found to be downregulated in tumor tissue compared to normal tissues. On the contrary, in tissues or fluids which contain low Vc levels, such as plasma and thyroid(Padayatty and Levine, 2016), the expression of SVCT2 in tumors was upregulated compared to normal tissues (Figure 2b). Thus, we hypothesized that the negative correlation between expression changes of SVCT2 in tumors and Vc concentrations in related tissues may be a regulatory mechanism for tumor cells to maintain ERK activity through the Vc signaling pathway mediated by SVCT2.

**Figure 2.**
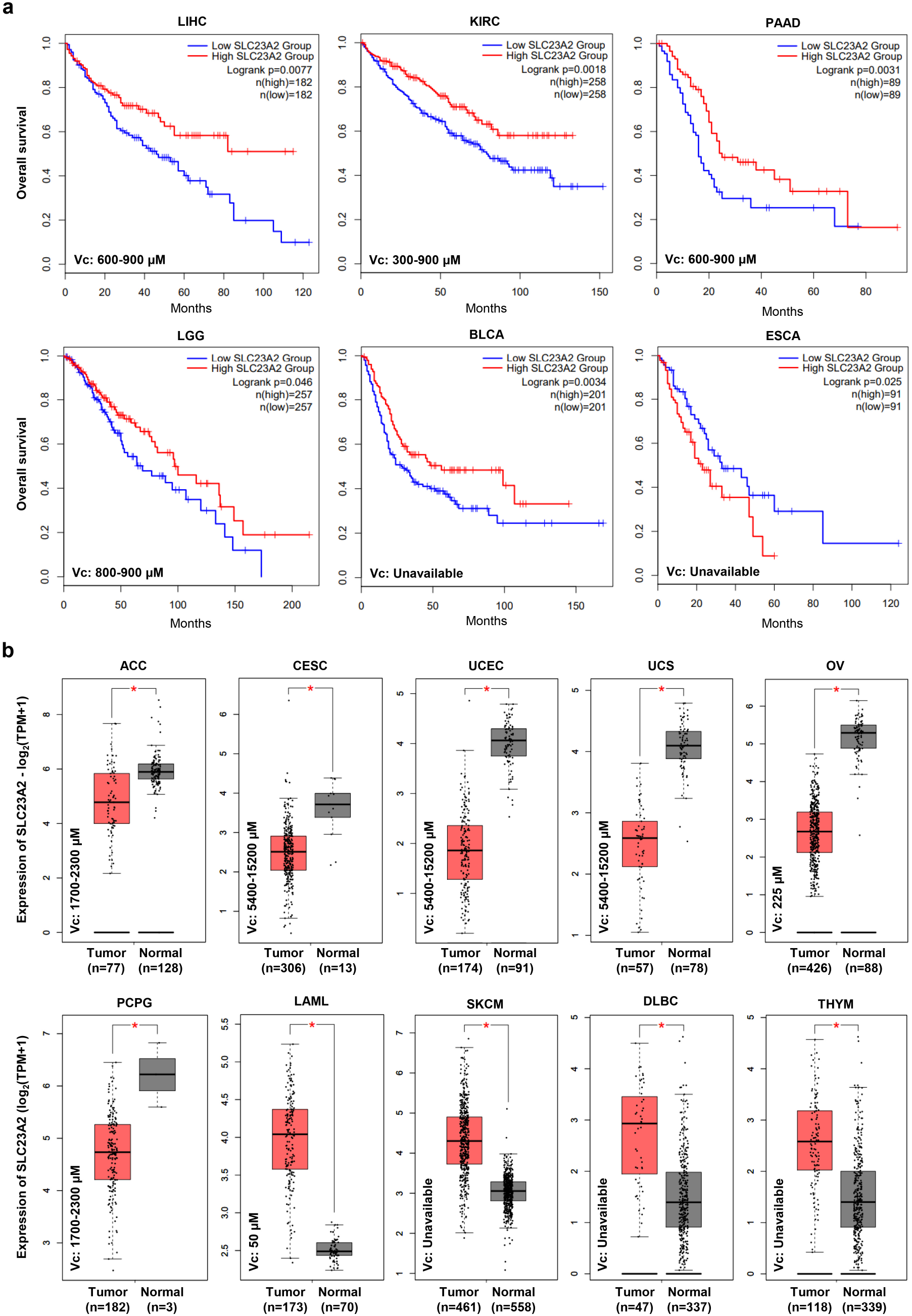
Survival and expression analysis of SVCT2 (SLC23A2) in various cancers. Survival (**a**) and differential expression (**b**) analysis of SVCT2 (SLC23A2) in various cancers were performed using GEPIA2 web server (http://gepia2.cancer-pku.cn/#index), and the expression and clinical features were from the TCGA and the GTEx projects. Lower left of each panel shows Vc concentrations in corresponding tissues or fluids. In BLCA, Vc concentrations in bladder is not reported, but the Vc concentration in urine is 200 μM as reported by Padayatty and Levine. In CESC, UCEC and UCS, exfoliated cervicovaginal cell Vc level was fourfold greater than that of leukocytes (1350-3800 μM) as reported by Basu et al. In OV, Vc concentration ratio is about 4.5:1 between follicular fluid and serum (50 μM), and ovary has been recognized as a site of Vc accumulation and turnover as reported by Luck et al. Other data were obtained from publication by Padayatty and Levine.

### Extracellular Vc binding to SVCT2 is sufficient for Vc-induced ERK activation

Due to the dual functions of mediating Vc uptake and signaling, phloretin (a transmembrane transport inhibitor of Vc) was applied to disrupt Vc transmembrane transport to reveal the relationship between Vc transmembrane transport and ERK signaling activity mediated by SVCT2 in HEK293T and F9 cells. Surprisingly, the activation of ERK was observed in all groups with Vc supplementation, irrespective of Vc transport inhibition (Figure 3a and S3a). Furthermore, wild-type SVCT2 and SVCT2 H109Q (a SVCT2 mutant without the ability for Vc transmembrane transport(Han et al., 2021; Ormazabal et al., 2010)) were overexpressed in HEK293T cells and F9 cells. Then the activation of ERK was detected after Vc stimulation. As expected, both SVCT2 H109Q and wild-type SVCT2 induced ERK activation in the presence of Vc (Figure 3b and S3b). These results confirmed that Vc-mediated activation of ERK was independent of Vc transmembrane transport.

**Figure 3.**
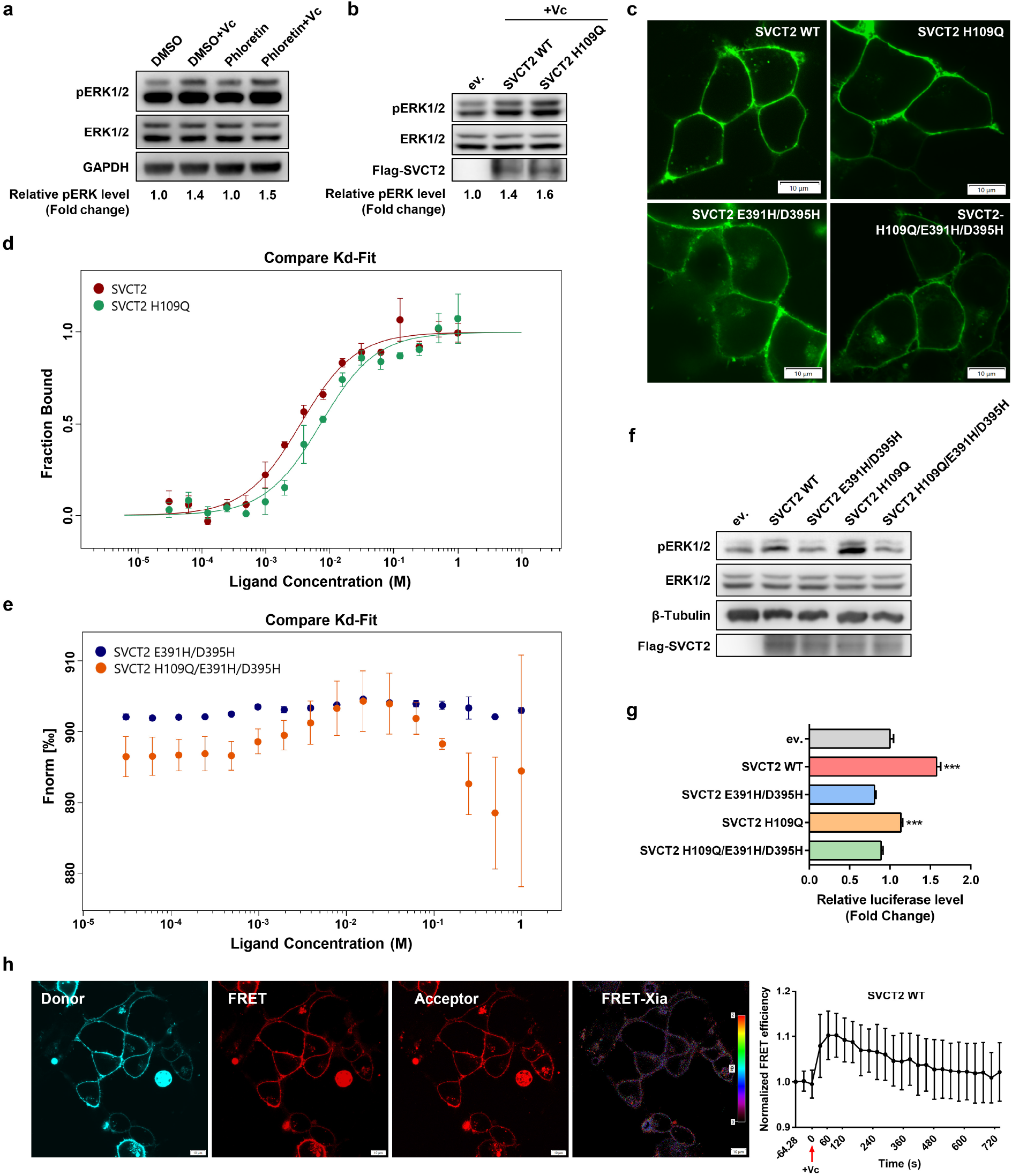
SVCT2 is a receptor mediating Vc-induced ERK activation. **(a)** HEK293T cells were pretreated with phloretin (200 μM) or DMSO for 4 h, then cells were treated with Vc (50 μg/mL) for 20 min, and lysates were immunoblotted as indicated. **(b)** HEK293T cells were transfected as indicated for 36 h and then treated with Vc (50 μg/mL) for 20 min. Lysates were immunoblotted as indicated. ev., empty vector. **(c)** HEK293T cells were transfected with GFP-fused wild-type SVCT2 (SVCT2 WT) or SVCT2 mutants for 36 h, then cells were imaged using confocal microscope. Scale bars, 10 μm. **(d and e)** Dose-response curve for the binding interaction between SVCT2/SVCT2 mutants and Vc was generated by MST analysis (Kd model). The concentration of GFP-fused SVCT2/SVCT2 mutants in HEK293T lysate was constant, the concentration of the L-ascorbate sodium was between 1000 mM to 0.0305 mM. Ligand concentrations on the x-axis were plotted in mole (M). **(f)** HEK293T cells were transfected as indicated for 36 h and then treated with Vc (500 μg/mL) for 2 min. Lysates were immunoblotted as indicated. ev., empty vector. **(g)** DLR assay of HEK293T cells co-transfected with SRE-luc pathway-profiling vectors and indicated vectors. 12 h after transfection, cells were incubated with Vc (50 μg/mL) and lysed 24 h later. **(h)** Time-Resolved FRET in living cells. Left: Representative confocal images of SVCT2-Clover (donor), SVCT2-mRuby2 (acceptor), FRET and FRET map. HEK293T cells were co-transfected with SVCT2-Clover and SVCT2-mRuby2 for 36 h, then cells were photographed using confocal microscope for sensitized emission FRET analysis. The FRET map resulting from the Xia method was displayed as pseudo color table. High values were displayed in red and low ratio values were displayed in magenta. Scale bar, 10 μm. Right: Time-scale FRET efficiency changes. For one time-scale, cells were treated with Vc (500 μg/mL) immediately after 3 pictures were snapped. Data in **h** were mean ± SD of the relative FRET efficiency (n=28 cells), and the maximum response after Vc treatment was significantly different to that of before Vc treatment (P<0.01, two-tailed Student’s *t*-test). Error bars represent SD of at least three independent experiments. ***, P<0.001; two-tailed Student’s *t*-test.

To further verify, SVCT2 E391H/D395H and SVCT2 H109Q/E391H/D395H mutants, which lacked the crucial docking site of Vc in the outward-facing gate of SVCT2(Kosti et al., 2012), were constructed. The immunofluorescence assay verified that GFP-tagged SVCT2 H109Q, SVCT2 E391H/D395H, SVCT2 H109Q/E391H/D395H, and wild-type SVCT2 were localized correctly to the membrane of HEK293T cells (Figure 3c). Then the binding of Vc to SVCT2 was further detected by microscale thermophoresis (MST). As shown in Figure 3d, a clear binding-induced change in MST was observed in both wild-type SVCT2 and SVCT2 H109Q, yielding a dissociation constant (*Kd*) of 3.46 mM in wild-type SVCT2 and 7.13 mM in SVCT2 H109Q with a right-shifted curve, while no significant binding-induced change in the MST signals was detected in SVCT2 E391H/D395H and SVCT2 H109Q/E391H/D395H mutants due to the loss of Vc-binding ability (Figure 3e). The results of western blot and DLR assay showed both SVCT2 E391H/D395H and SVCT2 H109Q/E391H/D395H were disabled to mediate ERK activation compared to wild-type SVCT2 and SVCT2 H109Q (Figure 3f and 3g). Moreover, ERK phosphorylation was induced by high-dose Vc in cells transfected with wild-type SVCT2 and SVCT2 H109Q but not in cells transfected with SVCT2 E391H/D395H and SVCT2 H109Q/E391H/D395H (Figure S3c). Collectively, these results suggested that the binding of Vc to the outward-facing of SVCT2 was sufficient for ERK activation and did not require transmembrane transport mediated by SVCT2, just as the canonical receptor-ligand binding.

### Transmembrane transport of Vc promotes dimerization of SVCT2

The oligomerization of transporters has been documented in previous studies. UapA and SVCT2 belong to the same family of nucleobase-ascorbate transporters (NATs), and it has been reported that UapA forms dimers on cell membranes(Alguel et al., 2016; Martzoukou et al., 2015). In order to reveal whether SVCT2 formed dimers on the cell membrane, SVCT2-mRuby and SVCT2-Clover were constructed for the Förster resonance energy transfer (FRET) assay. Sensitized emission FRET showed that FRET signals were detectable at the cell surface, suggesting SVCT2 may forms dimers on the cell membrane (Figure 3h). When ligands were bound to their cell surface receptor, a conformational change was triggered to facilitate receptor dimerization. Next, dynamic structural changes in the intracellular segment of SVCT2 induced by Vc were monitored by Time-Resolved FRET (TR-FRET) in living cells. TR-FRET revealed that Vc (500 μg/mL) stimulation increased the association between SVCT2-mRuby and SVCT2-Clover within 2 min, and then the FRET efficiency reduced gradually (Figure 3h), indicating that Vc induced the proximity of SVCT2. However, both SVCT2-H109Q and inhibition of Vc transport by phloretin resulted in the elimination of Vc-induced FRET efficiency increases (Figure S3d), suggesting that the dimeric form of SVCT2 was not necessary for ERK activation. Meanwhile, Tg (Tg101348 or Fedratinib, a specific inhibitor of JAK2), which significantly inhibited the transmembrane transport of Vc, also inhibited Vc-induced FRET efficiency increases (Figure S3d). Suggesting that the dimeric form of SVCT2 induced by Vc may depend on the transmembrane transport of Vc.

To investigate whether ERK activity was required for Vc transmembrane transport or Vc binding in turn, HPLC and MST assays were performed. As shown in Figure S3e, the intracellular accumulation of Vc in F9 cells with PD (PD0325901 or Mirdametinib, a specific inhibitor of MEK/ERK pathway) treatment decreased slightly compared to cells without PD treatment (Figure S3e). The intracellular Vc concentration was also detected under different doses of Vc supplementation using HPLC. The uptake of Vc increased gradually and tended to be steady, from 50 μg/mL of Vc along with ERK activation to 500 μg/mL without ERK activation (Figure S3f). MST confirmed that the inhibition of ERK exerted no effect on Vc binding to SVCT2 with a *Kd* of 3.45 mM, which was comparable to that without ERK inhibition (Figure S3g). Furthermore, PD treatment had no influence on Vc-induced FRET efficiency increases (Figure S3d). These findings suggested that ERK activity induced by Vc had no direct bearing on binding of Vc to SVCT2, the dimerization of SVCT2 and the Vc transport.

### The JAK2-GRB2 axis contributes to Vc-induced ERK activation

Till now, we have shown that SVCT2 mediated Vc-induced activation of JAK2 and ERK. Next, we wondered whether there was an interplay between Vc-induced activation of ERK and JAK2. To validate this hypothesis, JAK2 inhibitor (Tg) was applied with or without Vc stimulation in F9 cells and HEK293T cells. Tg treatment suppressed both basal and Vc-induced phosphorylation of ERK (Figure 4a and S4a), suggesting that JAK2 activity was required for Vc-induced ERK activation.

**Figure 4.**
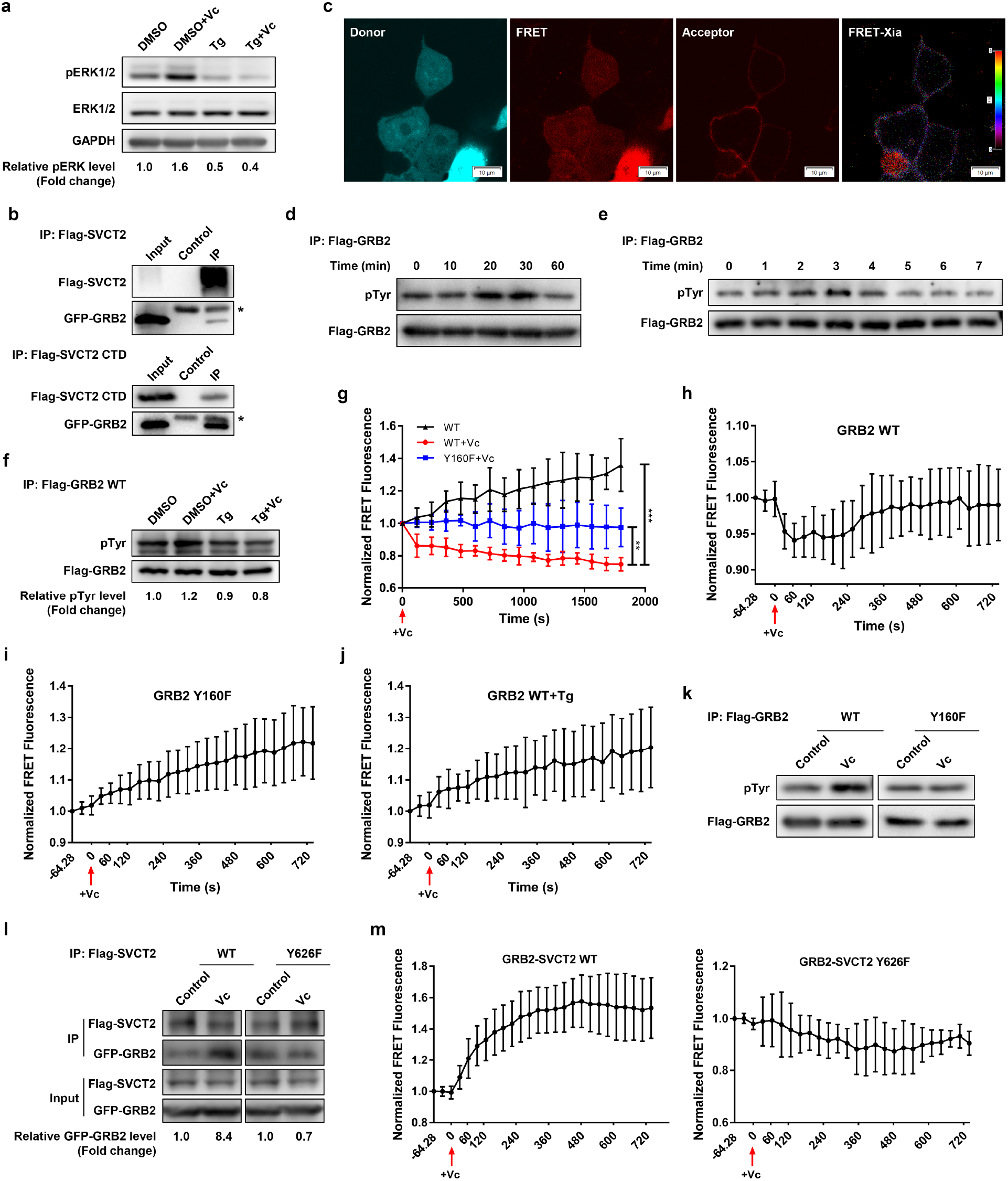
The JAK2-GRB2 axis contribute to Vc-induced ERK activation. **(a)** F9 cells were pretreated with Tg (2 μM) or DMSO for 4 h. After pretreatment, cells were treated with Vc (50 μg/mL) for 20 min. Lysates were immunoblotted as indicated. **(b)** HEK293T cells were co-transfected with Flag-tagged SVCT2 or SVCT2 CTD and GFP-GRB2 for 36 h, Flag-tagged SVCT2 or SVCT2 CTD was immunoprecipitated respectively and precipitates were detected for GFP-tagged GRB2. *, heavy chain of antibody. Control, pCDNA3.1(+) fused with triple Flag tag were co-transfected with GFP-GRB2. Flag-tag was immunoprecipitated and precipitates were detected for GFP-tagged GRB2. **(c)** Representative confocal images of GRB2-clover (donor), SVCT2-mRuby2 (acceptor), FRET and FRET map. HEK293T cells were co-transfected with GRB2-clover and SVCT2-mRuby2 for 36 h, then cells were photographed using confocal microscope for sensitized emission FRET analysis. The FRET map resulting from the Xia method was displayed as pseudo color table. High values were displayed in red and low ratio values were displayed in magenta. Scale bar, 10 μm. **(d and e)** HEK293T cells were transfected with Flag-tagged GRB2 for 36 h, then cells were treated with 50 μg/mL Vc (**d**) or 500 μg/mL Vc (**e**) and lysed at indicated times. Flag-tagged GRB2 was immunoprecipitated, and precipitates were blotted for phospho-tyrosine. pTyr, phospho-tyrosine. **(f)** HEK293T cells were transfected with Flag-tagged GRB2 for 36 h, cells were pretreated with Tg (2 μM) or DMSO for 4 h, then cells were treated with Vc (50 μg/mL) for 20 min. Flag-tagged GRB2 was immunoprecipitated, and precipitates were blotted for phospho-tyrosine. **(g)** Time-Resolved FRET in living cells. HEK293T cells were co-transfected with GRB2-Clover and GRB2-mRuby2 or GRB2 Y160F-Clover and GRB2 Y160F-mRuby2 for 36 h, Vc (50 μg/mL) was supplied to cells as indicated and cells were photographed using confocal microscope immediately for FRET analysis. Representative confocal images of donor, FRET, acceptor and FRET map were exhibited in Figure S4**e**. **, P<0.01, ***, P<0.001, two-tailed Student’s *t*-test, data in the last time point was analyzed. **(h-j and m)** Time-Resolved FRET in living cells. HEK293T cells were co-transfected with GRB2-Clover and GRB2-mRuby2 (**h** and **j**), GRB2 Y160F-Clover and GRB2 Y160F-mRuby2 (**i**), GRB2-Clover and SVCT2-mRuby2 or GRB2-Clover and SVCT2 Y626F-mRuby2 (**m**) respectively. 36 h after transfection, cells were treated with Tg (2 μM) (**j**) or DMSO (**h** and **i**) as indicated for 4 h and photographed time stack images using confocal microscope for sensitized emission FRET analysis. For one time series, cells were treated with Vc (500 μg/mL) immediately after 3 pictures were snapped. In **h** and the left panel in **m**, the maximum response after Vc treatment was significantly different to that of before Vc treatment (P<0.01, two-tailed Student’s *t*-test). **(k)** HEK293T cells were transfected with Flag-tagged wild-type GRB2 or GRB2 Y160F for 36 h, then cells were treated with Vc (50 μg/mL) for 20 min and lysed immediately. Flag-tagged GRB2 was immunoprecipitated, and precipitates were blotted for phospho-tyrosine. **(l)** HEK293T cells were co-transfected with Flag-tagged SVCT2 WT/SVCT2 Y626F and GFP-GRB2 for 36 h, then cells were treated with Vc (500 μg/mL) for 2 min. Flag-SVCT2/SVCT2 Y626F was immunoprecipitated and precipitates were blotted for GFP-GRB2. All data in **g**-**j** and **m** were mean ± SD of the relative FRET efficiency.

The adaptor growth factor receptor-bound protein 2 (GRB2) provided a critical bridge between cell surface receptors and the Ras signaling pathway(Ahmed et al., 2015; Haines et al., 2009). To understand whether GRB2 linked SVCT2 to the ERK pathway, immunofluorescence, co-Immunoprecipitation (Co-IP) and FRET experiments were performed to determine the interaction between GRB2 and SVCT2. As shown in Figure S4b, the immunofluorescence results showed that GFP-GRB2 and Flag-SVCT2 colocalized in the cell membrane. Co-IP experiments indicated that GRB2 was specifically associated with SVCT2 but not SVCT1 (Figure 4b and S4c), and the C-terminal of SVCT2 (SVCT2 CTD), an intracellular segment of SVCT2 which had previously been proven to be critical in JAK2/STAT2 signaling pathway, could recruit GRB2 (Figure 4b and S4d). FRET analysis finally revealed that SVCT2 interacted with GRB2 on the cell membrane (Figure 4c).

Previous studies have shown that phosphorylation of GRB2 by JAK2 acts as a bridge in connecting JAK2 and RAS/MEK/ERK pathway(Haines et al., 2009; Riera et al., 2010). In the present study, Flag-GRB2 overexpressed in HEK293T cells was immunoprecipitated (IP), and phospho-tyrosine (pTyr) was detected. The pTyr levels of GRB2 increased significantly at 20-30 min after Vc stimulus and decreased at 60 min (Figure 4d), consistent with the fluctuating trend of JAK2 activation induced by Vc(Han et al., 2021). Similarly, with high-dose Vc (500 μg/mL) stimulation, the level of pTyr on GRB2 increased at 2-3 min and then decreased after 4 min (Figure 4e). Furthermore, JAK2 inhibition by Tg abrogated the pTyr level of GRB2 induced by Vc (Figure 4f), suggesting JAK2 activity contributed to GRB2 phosphorylation.

As reported, only monomeric GRB2 is functional in upregulating RAS/MEK/ERK signaling transduction while the dimeric state is inhibitory to this process(Ahmed et al., 2015). To validate whether Vc promoted the dissociation of GRB2 dimers, HA-GRB2 and Flag-GRB2 were co-expressed in HEK293T cells. As shown in Figure S4f, the association between GRB2 was disrupted after Vc stimulation. The dynamic changes of GRB2 dimers under Vc stimulation also validated this result. TR-FRET revealed that GRB2 dimer was dissociated after Vc stimulation, which was observed at low (50 μg/mL) and high-dose (500 μg/mL) Vc stimulation (Figure 4g, S4e and 4h).

The conversion of GRB2 from a dimeric state to a monomer form has been reported to be facilitated by phosphorylation of GRB2 at Tyr^160^, which is located at the dimer interface and could be phosphorylated by JAK2(Ahmed et al., 2015; Haines et al., 2009; Riera et al., 2010). Hence, we sought to determine whether the phosphorylation of GRB2 at Tyr^160^ was related to JAK2 activation induced by Vc. TR-FRET showed that both the disruptive Tyr-Phe substitution of GRB2 at Tyr^160^ and JAK2 inhibitor Tg abrogated the dissociation of GRB2 dimer induced by Vc (Figure 4i and 4j), suggesting that the dissociation of the GRB2 dimer induced by Vc was associated with phosphorylation of GRB2 at Tyr^160^ and JAK2 activity. The pTyr levels of GRB2 in response to Vc was detected, results showed that the tyrosine phosphorylation increased on wild-type GRB2 after Vc stimulation and was not present in GRB2 Y160F (Figure 4k).

Tyrosine phosphate-containing ligand can serve as a recognition site for the SH2 domain of monomeric GRB2 after receptor activation. Hence, we wondered whether the phosphorylation of SVCT2 at Tyr^626^ induced by Vc-JAK2 reported previously(Han et al., 2021) could be a potential docking site for GRB2. As shown in Figure 4l and S4g, both wild-type SVCT2 and SVCT2 Y626F mutant could interact with GRB2; however, Vc increased the association of GRB2 with wild-type SVCT2 rather than SVCT2 Y626F. Similarly, TR-FRET showed that the association between GRB2 and SVCT2 was significantly increased within 6 min after Vc supplementation, while the association between GRB2 and SVCT2 Y626F no longer responded to Vc stimulation (Figure 4m), indicating that SVCT2 Tyr^626^ was related to recruitment of GRB2. Collectively, these results revealed that the activation of JAK2 induced by Vc promoted GRB2 monomerization through phosphorylating GRB2 and facilitated GRB2 recruitment to the cell membrane by phosphorylating SVCT2, thereby promoting ERK activation synergically.

### Vc-induced ERK activation phosphorylates PTPN12 at Ser^434^ to inhibit its phosphatase activity

To determine whether ERK activity was required for JAK2 activation induced by Vc, the phosphorylation of JAK2 at Tyr^1007/1008^ (pJAK2) was detected under PD treatment. The results showed that treatment with PD significantly abrogated JAK2 phosphorylation induced by Vc (Figure 5a), suggesting that ERK activity contributed to JAK2 activation. To decipher the roles of ERK in JAK2 activation induced by Vc, a quantitative phosphoproteome of Vc-stimulated F9 cells was performed. Phosphoproteome data revealed that phosphorylation of PTPN12 at Ser^434^ was significantly upregulated after Vc treatment (Figure S5a). Next, serine phosphorylation (pSer) of PTPN12 was validated with Vc stimulation over time. As shown in Figure 5b, the pSer level of PTPN12 peaked at 20 min after Vc (50 μg/mL) supplementation and then began to decrease after 30 min, which was consistent with the fluctuating trend of ERK phosphorylation induced by Vc. Additionally, Vc supplementation enhanced the pSer of PTPN12, whereas PD almost abolished the influence of Vc on inducing pSer of PTPN12 (Figure 5c). The pSer level of PTPN12 under both low and high-dose Vc in HEK293T cells and HeLa cells was also detected. As expected, the pSer levels of PTPN12 still increased under low-dose Vc stimulation in both cells, while there were no significant changes in the pSer levels of PTPN12 in HEK293T cells and a slight decrease in HeLa cells after being treated with high-dose Vc (Figure S5b). These results were consistent with ERK activity in both cells under low or high-dose Vc treatment (Figure 1m and S8e). All the results suggested that Vc-triggered ERK activation resulted in the serine phosphorylation of PTPN12.

**Figure 5.**
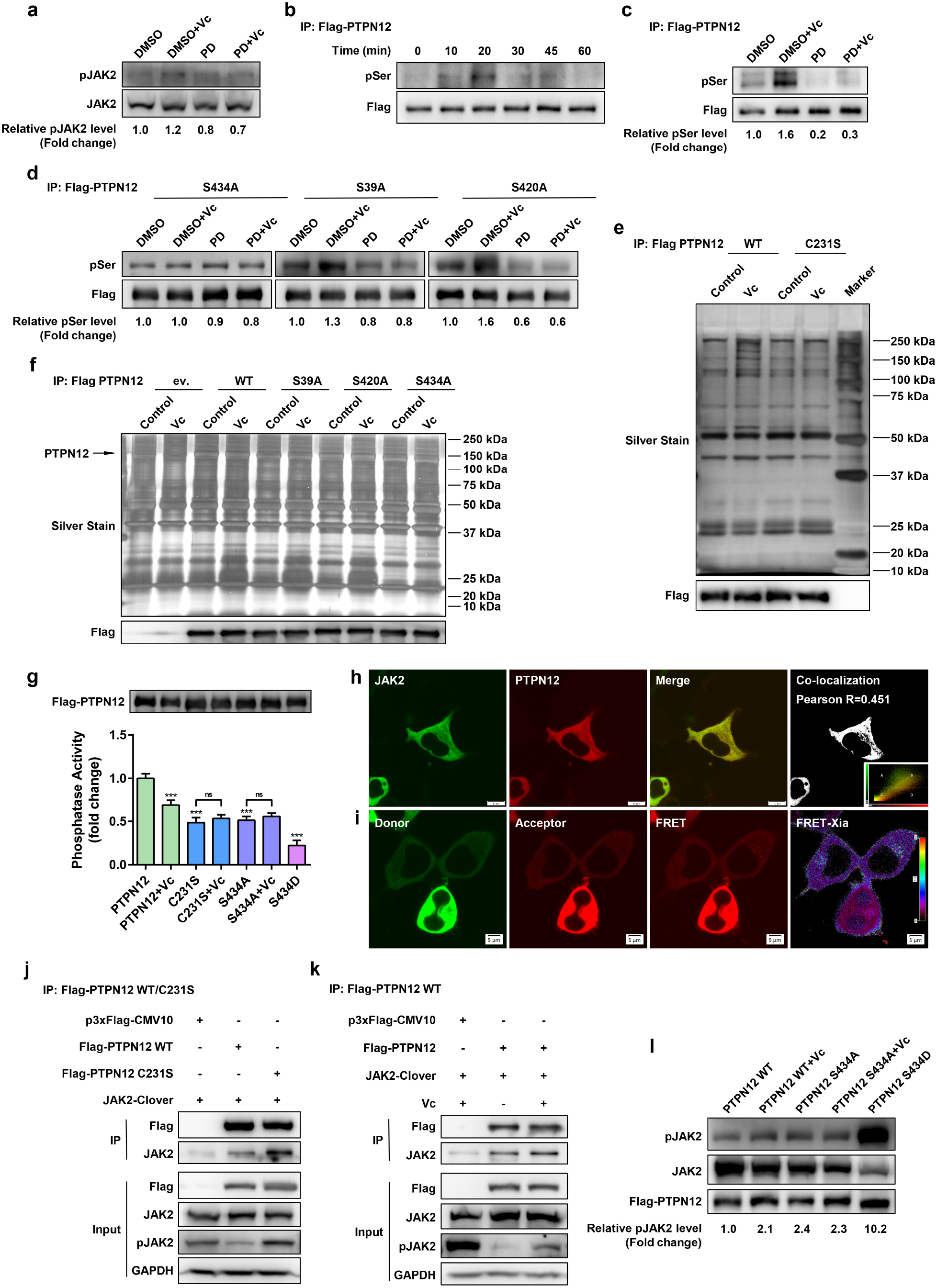
Vc-induced ERK activation phosphorylates PTPN12 at Ser^434^ to inhibit its phosphatase activity. **(a)** F9 cells were pretreated with PD (2 μM) or DMSO for 4 h, then cells were treated with Vc (50 μg/mL) for 30 min. Lysates were immunoblotted as indicated. The effect of PD was validated in Figure S5**c**. **(b)** HEK293T cells were transfected with Flag-tagged PTPN12 for 36 h, cells were treated with Vc (50 μg/mL) for indicated times. Flag-tagged PTPN12 was immunoprecipitated, and precipitates were blotted for phospho-serine. pSer, phospho-serine. **(c and d)** HEK293T cells were transfected with Flag-tagged PTPN12 (**c**) or PTPN12 mutants (**d**) as indicated for 36 h, cells were pretreated with PD (2 μM) or DMSO for 4 h. Then cells were treated with Vc (50 μg/mL) for 20 min and lysed immediately. Flag-tagged PTPN12 was immunoprecipitated, and precipitates were blotted for phospho-serine. **(e)** HEK293T cells were transfected with Flag-tagged wild-type (WT) PTPN12 or PTPN12 C231S for 36 h, cells were treated with Vc (50 μg/mL) for 20 min and lysed immediately. Flag-tagged PTPN12 was immunoprecipitated, and precipitates were analyzed by silver stain. **(f)** HEK293T cells were transfected with Flag-tagged wild-type PTPN12, PTPN12 mutants or empty vector (ev.) respectively as indicated for 36 h, then cells were treated with Vc (50 μg/mL) for 20 min and lysed immediately. Flag-tagged PTPN12 was immunoprecipitated, and precipitates were analyzed by silver stain. **(g)** Evaluation of phosphatase activity of PTPN12 and PTPN12 mutants. HEK293T cells were transfected with Flag-tagged wild-type PTPN12 or PTPN12 mutants for 36 h, then cells were treated with Vc (50 μg/mL) for 20 min. Flag-tagged PTPN12 was immunoprecipitated, and precipitates were analyzed using pNPP as a substrate for phosphatase activity or blotted for Flag tag. Error bars represent SD of five replicates. ns, not significant. ***, P<0.001, two-tailed Student’s *t*-test. **(h)** The colocalization between Flag-PTPN12 and JAK2-Clover in HEK293T cells. The illustration on the lower right showed a scatterplot in which the pixels were located more or less along a diagonal line. All of the pixels whose intensity in the B quadrant were shown in white. Scale bar, 10 μm. **(i)** Representative confocal images of JAK2-Clover (donor), PTPN12-mRuby2 (acceptor), FRET and FRET map. HEK293T cells were co-transfected with JAK2-Clover and PTPN12-mRuby2 for 36 h, then cells were photographed using confocal microscope for sensitized emission FRET analysis. The FRET map resulting from the Xia method was displayed as pseudo color table. High values were displayed in red and low ratio values were displayed in magenta. Scale bar, 5 μm. **(j and k)** HEK293T cells were co-transfected with Flag-tagged PTPN12 WT/PTPN12 C231S and JAK2-Clover as indicated for 36 h, then cells were lysed and Flag-PTPN12 was immunoprecipitated. The precipitates and input were blotted as indicated. In **k**, cells were treated with Vc (50 μg/mL) for 20 min. **(l)** HEK293T cells were co-transfected with Flag-tagged wild-type PTPN12/PTPN12 mutants and JAK2-Clover as indicated for 36 h, then cells were treated with Vc (50 μg/mL) for 20 min and lysed immediately. Lysates were immunoblotted as indicated.

To validate whether the phosphorylation of PTPN12 at Ser^434^ is in response to Vc-induced ERK activation, we constructed PTPN12 mutant PTPN12 S434A, along with two control mutants: PTPN12 S39A (both Ser^39^ and Ser^434^ of PTPN12 can be phosphorylated by PKA in previous reports(Garton and Tonks, 1994) and PTPN12 S420A (Ser^420^ is a predicted ERK phosphorylation site). Immunofluorescence assay validated that the mutation of PTPN12 at Ser^39^, Ser^420^ and Ser^434^ did not change the subcellular localization of PTPN12 (Figure S5e). Results of the IP assay showed that Vc promoted the pSer levels of PTPN12 S39A and PTPN12 S420A, which were significantly eliminated with PD treatment. However, pSer levels of PTPN12 S434A showed no response to Vc and PD (Figure 5d), suggesting Ser^434^ may be the main phosphorylation site of ERK induced by Vc. Nevertheless, H 89 2HCl, a specific inhibitor of PKA, could not block the basal and Vc-induced PTPN12 phosphorylation (Figure S5d).

PTPN12 is a non-receptor protein tyrosine phosphatase (PTP) family member with a deep pocket for phospho-tyrosine recognition(Andersen et al., 2001). It has been demonstrated that PTP mutants did not possess catalytic activity but still appeared to bind tightly to tyrosine-phosphorylated substrates(Jia et al., 1995). We adopted the substrate-trapping method to test the effect of Vc on the activity of PTPN12. Wild-type PTPN12 and PTPN12 C231S (an inactive mutant without the key catalytic amino acid cysteine) were immunoprecipitated from HEK293T cell lysate, and precipitates were analyzed by SDS-PAGE and silver staining. Wild-type PTPN12 pulled down more proteins in response to Vc treatment, while PTPN12 C231S did not respond to Vc stimulation (Figure 5e). In addition, the levels of pTyr in the precipitates were also detected. As shown in Figure S5f, Vc treatment resulted in more tyrosine phosphorylation in the proteins co-precipitated with wild-type PTPN12, while even more phosphorylated tyrosine contents that were not affected by Vc were detected in the proteins co-precipitated with the PTPN12 C231S inactive mutant, suggesting that Vc may induce inactivation of PTPN12.

In addition, the levels of proteins that co-precipitated with wild-type PTPN12 as well as PTPN12 S39A and PTPN12 S420A mutants were more enriched from cells with Vc supplementation than from cells without Vc treatment. On the contrary, PTPN12 S434A displayed lower substrate affinity and almost lost the response to Vc stimulation (Figure 5f), suggesting that the phosphorylation of PTPN12 at Ser^434^ resulted in the inactivation of PTPN12. To further confirm the above results, PTPN12 S434D, a mimicking phosphorylation mutant, was constructed. Then, we performed phosphatase assays in vitro using pNPP (p-Nitrophenyl phosphate) as a phosphatase substrate. As shown in Figure 5g, for wild-type PTPN12, significantly decreased activity was observed under Vc stimulation, showing an almost ∼25% reduction compared to Vc-untreated cells. In contrast, the PTPN12 C231S mutant was accompanied by a reduction in phosphatase activity with no response to Vc. Although different from the results of the substrate trapping assay, PTPN12 S434A exhibited an unexpected decrease in phosphatase activity, which may be accounted for by the fact that pNPP is only a small chemical molecule rather than a protein. However, the phosphatase activity of PTPN12 S434A did not respond to Vc supplementation, while PTPN12 S434D showed even lower phosphatase activity, suggesting that phosphorylation of PTPN12 at Ser^434^ induced by Vc led to the inhibited activity of PTPN12. Collectively, the Vc-ERK pathway inhibited the phosphatase activity of PTPN12 by phosphorylating PTPN12 at Ser^434^.

### ERK-PTPN12 axis contributes to Vc-induced JAK2 activation

To determine whether PTPN12 acts as a bridge between ERK and JAK2, the subcellular localization of JAK2 and PTPN12 in HEK293T cells was examined. As expected, JAK2 was colocalized with PTPN12 in the cytoplasm and membrane (Figure 5h). Then, sensitized emission FRET and the Co-IP assay confirmed that JAK2 interacted with PTPN12 in HEK293T cells (Figure 5i and 5j). Moreover, as shown in Figure 5j, the phosphorylation of JAK2 in cells co-expressing wild-type PTPN12 was significantly lower than that in cells without exogenous PTPN12, while pJAK2 levels in cells co-expressing inactive mutant PTPN12 C231S were significantly increased, suggesting that JAK2 may be a substrate of PTPN12. The results of substrate trapping experiment validated this finding: JAK2 co-precipitatation with wild-type PTPN12 was significantly less than with PTPN12 C231S mutant.

To verify whether PTPN12 inactivation induced by Vc was associated with JAK2 activation, wild-type PTPN12 was co-expressed with JAK2 in HEK293T cells with or without Vc supplementation. Vc stimulation resulted in more JAK2 pulldown with PTPN12 (Figure 5k). Accordingly, PTPN12 significantly promoted JAK2 dephosphorylation in the absence of Vc, while JAK2 phosphorylation was significantly increased in the presence of Vc due to the decreased activity of PTPN12 (Figure 5k).

Consistent with the results above, significant phosphorylated JAK2 co-expression with wild-type PTPN12 was induced by Vc. However, phosphorylated JAK2 co-expressed with PTPN12 S434A did not respond to Vc. Moreover, the low phosphatase activity mutant PTPN12 S434D resulted in a dramatic increase in JAK2 phosphorylation (Figure 5l). These results indicated Vc-induced ERK activation phosphorylated PTPN12 at Ser^434^ and inhibited its activity, then attenuated the dephosphorylation of JAK2.

Taken together, our findings revealed that Vc triggers an ERK-centered positive feedback loop consisting of ERK, PTPN12, JAK2 and GRB2, through its receptor-like transporter SVCT2, to maintain the activation of ERK.

### *Ptpn12* knockdown affects the progression of cancer

Based on the finding that the tumor suppressor PTPN12 is a downstream substrate of ERK induced by Vc, we hypothesized that PTPN12 might mediate the phenotype regulation of cancer cells by Vc. To substantiate that the ERK-PTPN12-JAK2 signaling axis also worked in cancer cells, *Ptpn12* was knockdown using the CRISPR/Cas9 system in a murine melanoma cell line B16-F10 (Figure S6a and S6b). The pJAK2 levels in *Ptpn12* knockdown cells were consistently higher than in wild-type B16-F10 cells (Figure 6a), supporting the above results that PTPN12 led to JAK2 dephosphorylation. Then, the phosphorylation of JAK2 and ERK in response to different doses of Vc in *Ptpn12* knockdown cells was analyzed. ERK phosphorylation in *Ptpn12* knockdown cells in response to Vc was found to be similar to the results obtained in wild-type cells, while the activation of JAK2 in *Ptpn12* knockdown cells showed almost no response to Vc (Figure 6b and 6c), suggesting PTPN12 indeed worked as a downstream substrate of ERK and upstream mediator of JAK2 in the positive feedback loop. Next, the phenotype of *Ptpn12* knockdown cells was examined. The results showed that knockdown of *Ptpn12* promoted the proliferation, migration and inhibited cell apoptosis of B16-F10 cells (Figure 6d-6f). Then xenograft tumor models using wild-type or *Ptpn12* knockdown B16-F10 cells were constructed. As expected, knockdown of *Ptpn12* promoted the growth of B16-F10 cell xenograft tumors (Figure 6g).

**Figure 6.**
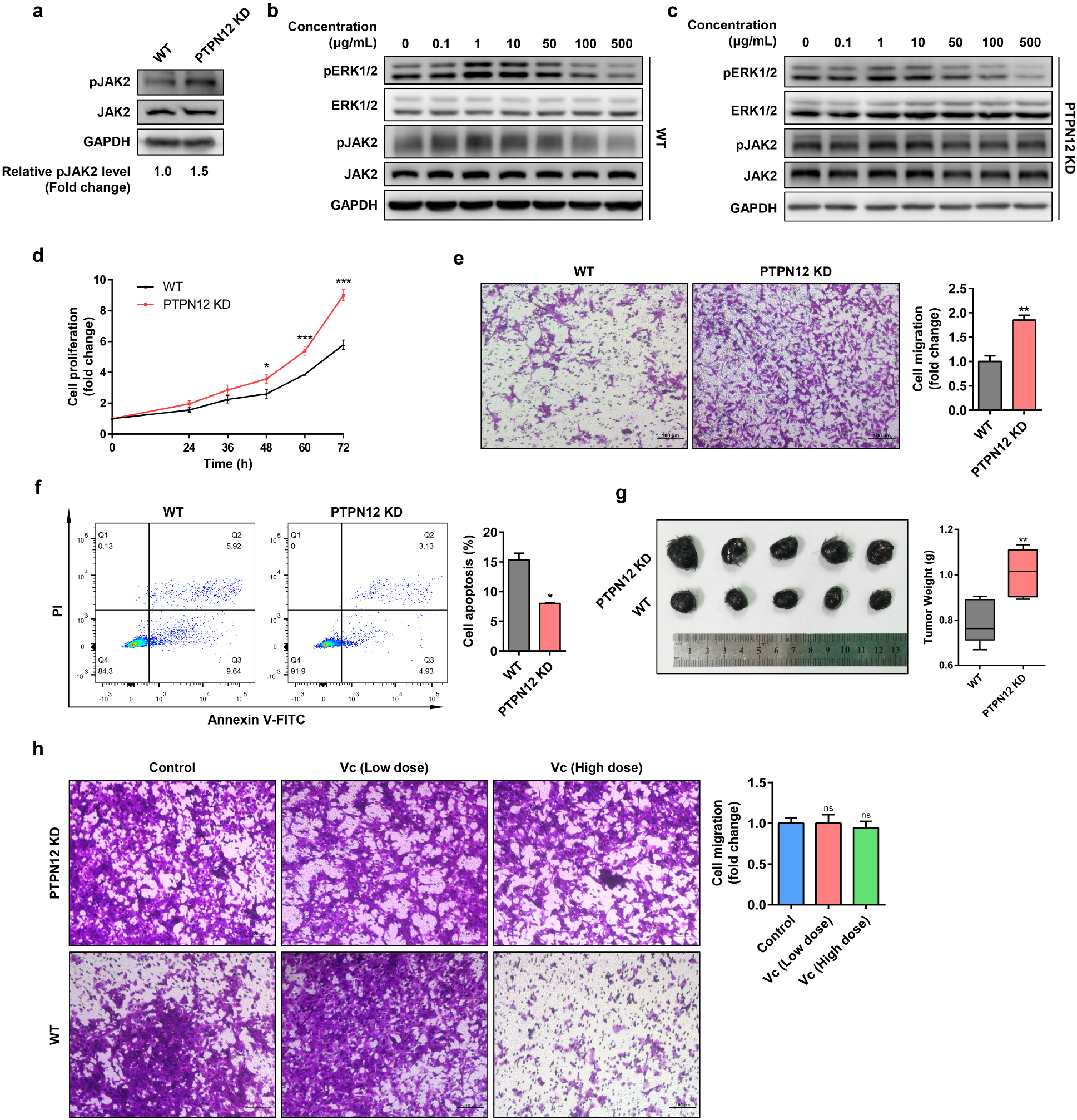
*Ptpn12* knockdown affects the progression of cancer. **(a)** Wild-type B16-F10 cells (WT) and *Ptpn12* knockdown B16-F10 cells (PTPN12 KD) were lysed and lysates were immunoblotted for endogenous pJAK2, total JAK2 and GAPDH. The knockdown efficiency was verified (data shown in Figure S6**a** and S6**b**). **(b and c)** Wild-type and *Ptpn12* knockdown B16-F10 cells were treated with Vc at indicated concentrations (0-500 μg/mL) for 20 min. Lysates were immunoblotted as indicated. **(d)** Cell proliferation of wild-type and *Ptpn12* knockdown B16-F10 cells were evaluated by CCK8 analysis. **(e)** The migration ability of wild-type and *Ptpn12* knockdown B16-F10 cells were detected by transwell assay. Scale bar, 100 μm. See also in Figure S6**c**. **(f)** Cell apoptosis of wild-type and *Ptpn12* knockdown B16-F10 cells were detected by flow cytometry. Results were representative of one of three similar experiments. **(g)** Effects of *Ptpn12* knockdown on xenograft tumor growth in vivo. Left: representative images of tumors in mice models constructed by wild-type or *Ptpn12* knockdown B16-F10 cells (n=5). Right: tumor weight was measured after tumor excision. **(h)** The migration ability of wild-type and *Ptpn12* knockdown B16-F10 cells treated with different doses of Vc were determined by transwell assay. Low dose, 10 μg/mL. High dose, 500 μg/mL. Scale bar, 100 μm. The migration ability of wild-type B16-F10 cells share the same dataset with Figure 7b. For all graphs, data were presented as mean ± SD. ns, not significant; *, P<0.05; **, P<0.01; ***, P<0.001; two-tailed Student’s *t*-test compared to control.

To illustrate the role of Vc in the regulation of PTPN12 activity, the phenotype of *Ptpn12* knockdown cells was detected under low and high-dose Vc treatment. Intriguingly, the migration ability of *Ptpn12* knockdown cells showed no response to Vc treatment regardless of the dose of Vc (Figure 6h), while the proliferation or apoptosis of *Ptpn12* knockdown cells was significantly inhibited or promoted by high-dose Vc (Figure S6d and S6e), similar to our findings in wild-type B16-F10 cells (Figure 7a-7c). No obvious change was observed with low-dose Vc treatment. The above findings suggested that the effect of Vc on tumor migration was mainly through PTPN12, which is consistent with previous reports that PTPN12 inhibits tumor migration(Lee and Rhee, 2019).

**Figure 7.**
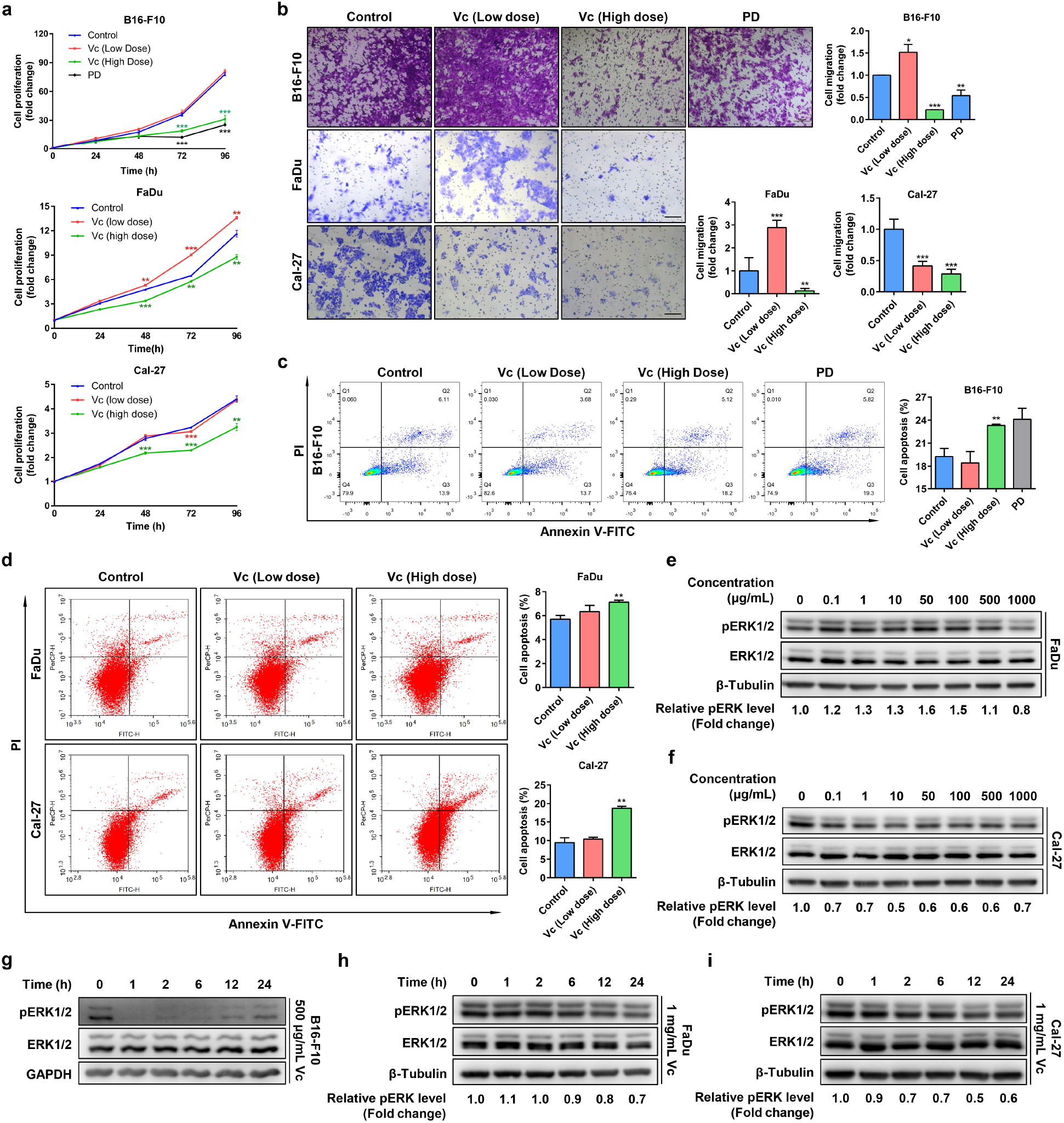
High-dose Vc affects the phenotype of cancer cells through inhibiting ERK. **(a)** Proliferation of B16-F10, FaDu and Cal-27 cells treated with different doses of Vc or PD (2 μM) were determined by CCK8 assays. **(b)** The migration ability of B16-F10, FaDu and Cal-27 cells treated with different doses of Vc, or combined PD (2 μM) were detected by transwell assays. Scale bar, 100 μm. See also Figure S7**j** and S7**k**. **(c and d)** Apoptosis of B16-F10, FaDu and Cal-27 cells treated with different doses of Vc were detected by flow cytometry. Results were representative of one of three similar experiments. **(e and f)** FaDu cells (**e**) and Cal-27 cells (**f**) were treated with Vc at indicated concentrations for 20 min. Lysates were immunoblotted as indicated. **(g, h and i)** B16-F10 cells (**g**), FaDu cells (**h**) and Cal-27 cells (**i**) were treated with Vc at indicated concentrations and lysed at indicated times. Lysates were immunoblotted as indicated. For B16-F10 cells, low dose was 10 μg/mL, high dose was 500 μg/mL; for FaDu and Cal-27 cells, low dose was 50 μg/mL, high dose was 1000 μg/mL. For all graphs, data were presented as mean ± SD. *, P<0.05; **, P<0.01; ***, P<0.001; two-tailed Student’s *t*-test compared to control.

### Vc regulates the progression of cancer through ERK

Due to the prevalence of ERK hyperactivation in cancers, we speculate that Vc affects cancer progression through ERK pathway mediated by SVCT2. Firstly, PD treatment, used as a positive control, showed that ERK inhibition significantly inhibited cell proliferation and migration ability, increased cell apoptosis in B16-F10 cells (Figure 7a-7c), which consistent with the classic perspective. Next, B16-F10 cell line, hypopharyngeal cancer cell line FaDu, human tongue squamous cell carcinoma cell line Cal-27 and human cervical adenocarcinoma cell line HeLa were treated with different doses of Vc. The results showed different phenotypic changes of these cancer cells with low-dose Vc treatment, especially in Cal-27 cells. The migration ability of B16-F10, Fadu and HeLa cells and the proliferation of Fadu cells were promoted with low-dose Vc. However, in Cal-27 cells, low-dose Vc inhibited the migration ability (Figure 7a, 7b, S8a and S8c). Conversely, similar to PD treatment, high-dose Vc significantly inhibited proliferation and migration ability and increased cell apoptosis in all cells (Figure 7a-7d and S8a-S8d).

Then, the changes of pERK levels in these cells were detected. The results revealed that the pERK levels in B16-F10, Fadu and HeLa cells were increased with low-dose Vc treatment and decreased with high-dose Vc treatment (Figure 6b, 7e and S8e). In contrast, pERK levels in both low and high-dose Vc-treated Cal-27 cells were decreased (Figure 7f), which fit the results perfectly that both low and high-doses Vc inhibited the migration ability of Cal-27 cells. Interestingly, according to the results in Figure 1k that increasing SVCT2 expression may resulted in ERK inhibition even under low-dose Vc stimulation, the result Cal-27 cells had the highest SVCT2 expression levels among these cancer cells (Figure S7h) may explain why ERK inhibition was found under both low and high-dose Vc treatment. Furthermore, pERK levels were further verified in cells treated with high or low-doses of Vc for different periods. The pERK levels in all high-dose Vc-treated cells and low-dose Vc-treated Cal-27 cells were decreased and lasted for at least 24 h (Figure. 7g-7i and S7c), on the contrary, the pERK levels were increased in B16-F10 and Fadu cells for at least 6 h after low-dose Vc stimulation (Figure S7a and S7b). These results suggested that the mechanism of Vc in affecting the phenotype of cancer cells could be explained by Vc-induced pERK level changes.

To corroborate that the above mechanisms found in HEK293T and F9 cells are also present in cancer cells, pERK levels in B16-F10, FaDu and HeLa cells treated with high-dose Vc also increased within 5 min and followed by feedback inhibition (Figure S7d, S7e and S8f). For Cal-27 cells, similar results could also be observed by increasing Vc concentration to 2000 μg/mL, with the activation time of ERK less than 1 min (Figure S7f and S7g), which could not be sampled in such a short lapse of time. In addition, the time-scale changes of pERK in response to high-dose Vc were not affected by the presence of phloretin in B16-F10 cells (Figure S7i).

We further constructed xenograft tumor models using B16-F10, FaDu and Cal-27 cells, and intratumoral injection (i.t.) with low or high-dose Vc was performed. Growth of B16-F10 and FaDu xenograft tumors was promoted by low-dose Vc but retarded by high-dose Vc (Figure 8a, 8b and S9a). In contrast, both low and high-doses of Vc significantly retarded the growth of Cal-27 xenograft tumors (Figure 8c and S9a). Immunohistochemical staining revealed that changes of pERK levels were consistent with tumor size, and Vc altered the levels of proliferation marker Ki67, and Epithelial-mesenchymal transformation (EMT) marker (E-cadherin, N-cadherin and Vimentin) in xenograft tumors (Figure 8d and S9b), which was consistent with results observed in vitro. Collectively, these findings suggested that Vc affected the proliferation, migration and tumorigenesis through the SVCT2-ERK-PTPN12-JAK2-GRB2-ERK positive feedback loop. Although high-dose Vc always leads to persistent ERK feedback inhibition, the ability to induce ERK inhibition in specific tumor cells seems to be the determining factor for the effective anti-tumor dose of Vc.

**Figure 8.**
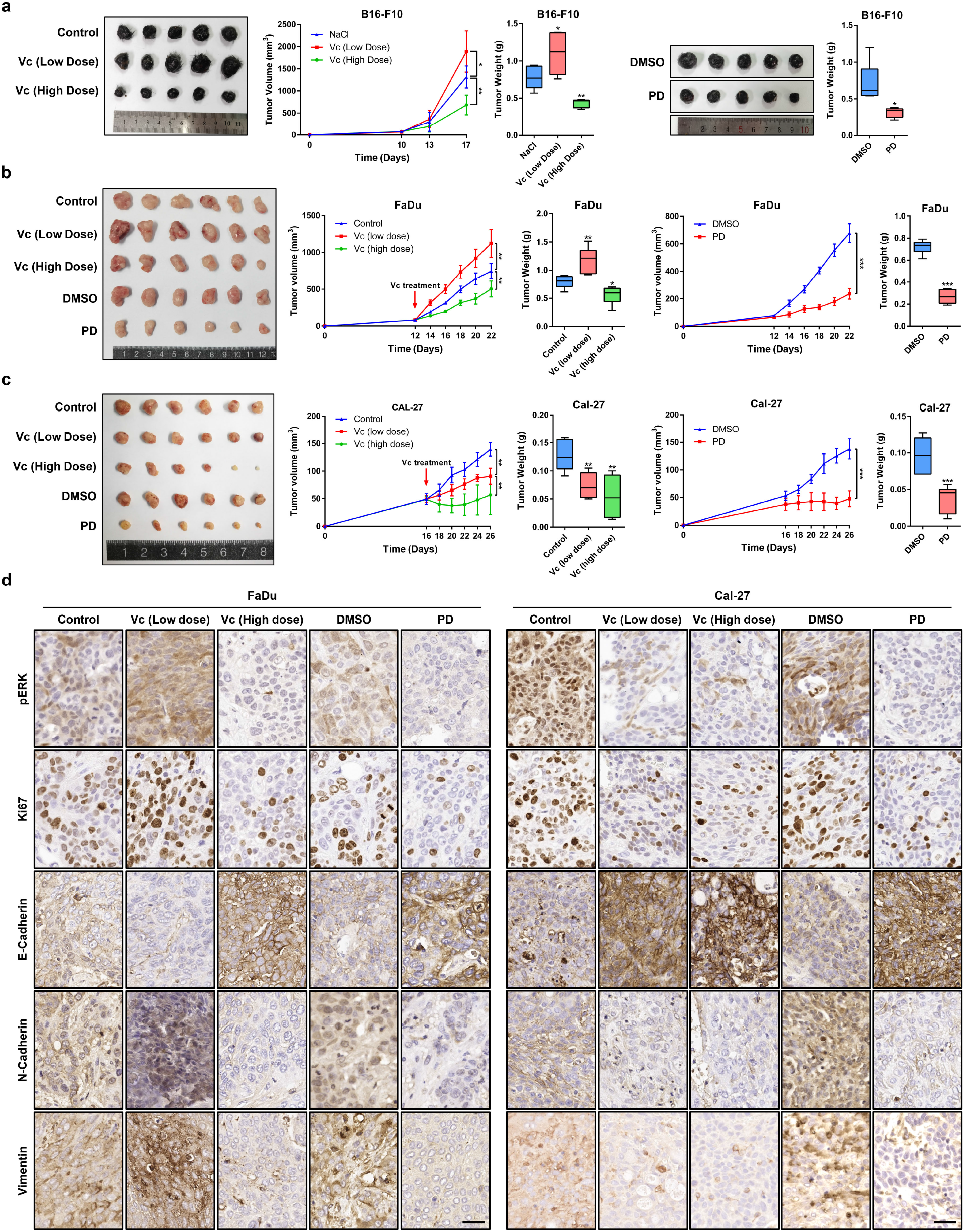
Vc affects tumorigenesis through the ERK pathway. **(a, b and c)** Effects of different doses of Vc and PD on xenograft tumor growth in vivo. Left: representative images of tumors in mice models constructed by B16-F10 cells (n=5), FaDu cells (n=6) and Cal-27 cells (n=6) after intratumoral injection of Vc or PD. Right: tumor growth curve was plotted using xenograft tumor volume data and tumor weight was measured after tumor excision. In DMSO and PD groups of B16-F10 xenograft tumor, the two images shared the same magnification. Data were presented as mean ± SD. *, P<0.05; **, P<0.01; ***, P<0.001; two-tailed Student’s *t*-test compared to control. **(d)** IHC staining of pERK, Ki67, E-Cadherin, N-Cadherin and Vimentin in xenograft tumors constructed by FaDu cells and Cal-27 cells. Scale bar, 40 μm.

## Discussion

Vc had also been found to exert other signaling-like functions except of inducing ERK activation, such as inhibiting NF-κB pathway through DHA(Bowie and ONeill, 2000; Cárcamo et al., 2004) and activating PKC through ROS generation(Baek et al., 2017). Since the explanation is almost perfect, and more importantly, no Vc receptor has been found, it is generally believed that Vc has no direct role in signal transduction(Smirnoff, 2018), even though several researches had proposed that Vc may not only acted as an antioxidant(Cárcamo et al., 2002; Esteban et al., 2010; Ulrich-Merzenich et al., 2007). Typically, the signal transduction pathway is initiated upon a signal binding to the extracellular domain of a cell surface receptor, without the need for the ligand to enter cells. Yun et al. reported that cancer cells hardly transport Vc(Yun et al., 2015), and our results demonstrated high-dose Vc still inhibited ERK after 10 min even though Vc transmembrane transport was blocked (Figure S7i). In addition, ERK activation induced by Vc could still occur when the Vc transmembrane transport was blocked by phloretin or transfected with SVCT2 H109Q mutant, but not in SVCT2 with E391H/D395H mutations (Figure 3a, 3b and 3f). These results indicated that binding of extracellular Vc to SVCT2 was sufficient for ERK activation and excluded all possible effects that may act on the ERK pathway induced by intracellular Vc and other effects induced by extracellular Vc. In this study, we provided further evidence to substantiate that SVCT2 functions as a receptor of Vc and Vc participates in intracellular signal transduction as a signaling molecule.

Activated receptors have been reported to provide a phosphorylated tyrosine as a docking site for GRB2, acting as a bridge between receptor tyrosine kinases (RTK) and the RAS/MAPK pathway, and hence resulting in SOS recruitment and activation of RAS(Buday and Downward, 1993). Previously, we found that Tyr^626^ of SVCT2, which was phosphorylated by JAK2, played a direct role in the recruitment of STAT2 to mediate the JAK2/STAT2 pathway(Han et al., 2021). In this study, we showed that the phosphorylated SVCT2 at Tyr^626^ could promote the association of GRB2 to SVCT2 and mediate activation of RAS-ERK cascade. In addition, significant Vc-induced ERK activation was observed several min following Vc stimulation, corroborating that the activation of ERK and JAK2 were not related to the intracellular accumulation of Vc and excluded the possibility that ERK activation was due to changes in gene expression induced by the roles of Vc in epigenetic regulation.

Dimerization is also essential for RTKs and tyrosine kinase-associated receptors to activate the tyrosine kinase activity by trans-autophosphorylation. We previously reported that Vc-induced JAK2 activation was tightly coupled with the transmembrane transport of Vc. However, the mechanism of JAK2 activation mediated by SVCT2 has not been elucidated. Here, combined with FRET assay, our results showed treatments that inhibited SVCT2 dimerization, such as SVCT2 H109Q mutant and phloretin or Tg, had been reported to disrupt Vc transmembrane transport and block JAK2 activation. Accordingly, we hypothesized that the dimerization of SVCT2 triggered by Vc transport could increase proximity between the two C-terminal and two JAK2, facilitating trans-autophosphorylation, then the activated JAK2 phosphorylated SVCT2 at Tyr^626^, which facilitated Vc transport in turn and mediated the activation of JAK2/STAT2 and RAS/ERK pathway by recruiting STAT2 and GRB2.

Vc-induced ERK activation was widespread in almost all cells detected, while significant JAK2 activation was only observed in F9 and B16 cells. Herein, we found intracellular PTPN12 expression level may be one of the reasons for the failure to observe significant Vc-induced phosphorylation of JAK2 at Tyr^1007/1008^ in most cells. As a tumor suppressor protein, the expression of PTPN12 is often low or undetectable in a variety of cancer cells(Meeusen and Janssens, 2018; Sun et al., 2011). We speculated that the insufficient dephosphorylation catalyzed by PTPN12 results in high basal level of phosphorylation on Tyr^1007/1008^ of JAK2 in these cells, which resulted in insensitive to Vc induction. As shown in this study, Vc-induced JAK2 phosphorylation was rescued in HEK293T cells with JAK2 and PTPN12 co-transfection (Figure 5l), in contrast to that with JAK2 transfection alone in previous paper. In addition, *Ptpn12* knockdown led JAK2 phosphorylation insensitive to Vc induction in contrast to wild-type B16-F10 cells (Figure 6b and 6c).

The ERK-PTPN12-JAK2-GRB2-ERK positive feedback loop can explain the mutual promotion between JAK2 and ERK activation induced by low and high-dose Vc for a short period. However, ERK inhibition under high-dose Vc treatment for a long period could not be explained by this mechanism. We speculate that this positive feedback loop tends to lead to ERK hyperactivation induced by Vc, so the cell adopts a negative feedback mechanism to fine-tune the activity of ERK for the purpose of maintaining cell homeostasis. As a result, significant oscillatory behavior of ERK phosphorylation induced by Vc was observed. And the long-term effects of high-dose Vc may be a result of this unknown negative feedback mechanism due to violent ERK activation induced by high-dose of Vc. Multiple negative feedback mechanisms targeting the ERK cascade have evolved, including direct posttranslational modification or regulation of the specific pathway inhibitors(Lake et al., 2016). Suggestively, the dephosphorylated members of this positive feedback loop, observed under high-dose Vc treatment, may be crucial targets of this feedback inhibition. Nevertheless, further studies are needed to elucidate the exact mechanism of this feedback inhibition.

High-dose Vc increases 5-hydroxymethylcytosine (5hmc) levels in cancer cells(Mikkelsen et al., 2021; Peng et al., 2018) and promotes H_2_O_2_ or ROS generation leading to DNA damage (Doskey et al., 2016; Su et al., 2019), are potential mechanisms of its anti-cancer effects. In contrast, our findings indicated that low-dose Vc exhibited anti-cancer effects on Cal-27 cells, which could not be explained by these two mechanisms. But the effects of Vc on ERK activity documented in this study could substantiate this observation. Notably, the sensitivity of different cells to Vc is not identical due to complicated signaling crosstalk. In several cancers, appropriate Vc intake may have a protective effect(Luo et al., 2014), and adequate dietary intake of Vc to maintain high levels of Vc in vivo may reduce the risk of certain types of cancer, such as squamous cell carcinoma of tongue.

The administration of Vc in cancer therapy through intravenous injection remains controversial. Human plasma Vc concentration was reported to be ∼50 μM, and Vc content inside the neutrophils can reach as high as 10 mM(Padayatty and Levine, 2016), thereby a pivotal question for Vc therapy is the effective dose. This study demonstrated that Vc acts as a signaling molecule to exert its roles in the ERK pathway, requiring direct interaction between Vc and SVCT2. Notwithstanding that lots of human phase I clinical trials achieved high-dose Vc in the blood through intravenous injections, no trials investigated the actual dose of Vc in contact with tumor cells. We assumed that the actual Vc concentrations reaching the tumor might be lower than the blood concentration. Accordingly, we attempted intratumoral injection of Vc, and our hypothesis was confirmed, as evidenced by the results of tumor progression and IHC. We also found that SVCT2 levels were associated with overall patient survival for some cancers, and expression changes of SVCT2 in tumors exhibited a negative correlation with Vc concentrations in corresponding tumor tissues. Based on our results, we concluded that both high-dose Vc and high expression of SVCT2 would lead to ERK inhibition. For LIHC, KIRC, PAAD, LGG and BLCA patients with high expression levels of SVCT2, we speculated that high doses of Vc in corresponding tumor tissues may lead to ERK inhibition, which in turn contributes to better prognosis. Conversely, for tumor self-maintenance, SVCT2 downregulation may be to avoid ERK inhibition in tumor tissues with high Vc concentration, while SVCT2 upregulation may be to maintain ERK at an activated state in tissues or fluids which containing low concentration of Vc. Accordingly, we believe that the response of ERK to Vc may be harnessed as a target for cancer therapy, and further studies are needed to quantify and optimize the concentrations of Vc reaching the target tissue. Our findings also suggest that some solid tumors could alternatively be treated with intratumoral injection of Vc instead of surgery.

In fact, we believe the word “transceptor” is more appropriate for SVCT2. It has been found that the transporters of some substances have receptor activity(Alguel et al., 2016; Martzoukou et al., 2015). So why cant SVCT2 be next? In conclusion, our study elucidated in detail the molecular mechanism of SVCT2 as a Vc receptor mediating the activation of the ERK-PTPN12-JAK2-GRB2-ERK positive feedback loop. Moreover, we found that this positive feedback loop existed widely in normal and cancer cells. In cancer cells, low-dose Vc activated the ERK pathway to promote malignant progression, while high-dose Vc resulted in feedback inhibition of the ERK pathway to suppress malignant phenotype. Therefore, our study provided new insights into the mechanism of Vc action and provided the basis for the development of new drug targets for cancer therapy. More importantly, our study provided new and important evidence for the application of Vc in cancer therapy.

## Acknowledgments

We thank Dr. Hua Zhao and Dr. Fengping Yuan (State Key Laboratory of Crop Stress Biology for Arid Areas, Northwest A&F University, Yangling, China) for confocal experimental assistance. We appreciate the Life Science Research Core Services (LSRCS, Northwest A&F University, Yangling, China). We also thank Menghao Qin, Nanjing University, for helpful discussions; and Yingbing Zhang, Northwest A&F University, for providing instructions of animal experiment. This work was supported by the National Natural Science Foundation of China (No. 31572405 to Z.G., No. 31872355 to Z.G., No. 32150002 to Z.G. and No. 82073101 to Y.W.), the Research Funds for China Central Government-guided Development of Local Science and Technology (No. 2020-165-19 to W.G.) and Shenzhen Key Laboratory Foundation (ZDSYS20200811143757022).

## Author contributions

Yian Guan, Bingxue Chen and Yongyan Wu contributed equally to this work.

Conceptualization: Yian Guan, Bingxue Chen, Zekun Guo; Methodology: Yian Guan, Bingxue Chen; Formal analysis: Yian Guan, Bingxue Chen, Yongyan Wu; Investigation: Yian Guan, Bingxue Chen, Yongyan Wu, Zhuo Han, Hongyu Xu, Caixia Zhang, Weijie Hao; Writing-original draft: Yian Guan, Bingxue Chen, Yongyan Wu, Zhuo Han, Zekun Guo; Writing-review & editing: Yian Guan, Bingxue Chen, Yongyan Wu, Zekun Guo; Funding acquisition: Zekun Guo, Yongyan Wu, Wei Gao; Resources: Wei Gao, Zekun Guo; Project administration: Wei Gao, Zekun Guo; Supervision: Wei Gao, Zekun Guo.

## Methods and materials

### Resource availability

#### Lead contact

Further information and requests for resources should be directed to the Lead Contact, Zekun Guo (E-mail address).

#### Materials availability

Plasmids and cell lines generated in this study are available on request from the Lead Contact.

#### Data and code availability

This paper does not report original code.

### Experimental model and subject details

#### Cells culture

All cell lines except Jurkat T lymphocyte leukemia cells were cultured in Dulbeccos modified Eagles medium (DMEM) (Thermo Fisher Scientific) supplemented with 10% FBS (Biological Industries) and 3.7 g/L NaHCO_3_ (Sigma-Aldrich) at 37°C, 5% CO_2_. Jurkat T lymphocyte leukemia cells were cultured in PRMI1640 (Thermo Fisher Scientific) supplemented with 10% FBS (Biological Industries) and 3.7 g/L NaHCO_3_ (Sigma-Aldrich) at 37°C, 5% CO_2_. All cells were tested free of mycoplasma contamination.

#### Animal models

B16-F10 xenograft tumor model: Animal protocols have been reviewed and approved by the Animal Ethical and Welfare Committee, Northwest A&F University, China. C57BL/6 mice were purchased from the Experimental Animal Center of Xi’an Jiaotong University (certificate No. SCXK [SHAAN] -2018-001). All mice were housed in isolated ventilated cages (maxima 6 mice per cage) at Northwest A&F University. Mice were maintained on a 12/12 h light (7:30-19:30) and dark cycle, 23 ± 2°C, 50 ± 10% humidity and given unrestricted access to standard diet and water. Five to six-week-old mice (C57BL/6) were used in the experiments. Littermates of the same sex (female) were randomly assigned to experiment groups. To construct xenograft tumor models, 50 μL sterilized normal saline containing 1×10^5^ B16-F10 cells were intracutaneously injected into the right flank of each mouse after induction of anesthesia.

FaDu/Cal-27 xenograft tumor model: Animal experiments were approved by the Animal Ethical and Welfare Committee, Northwest A&F University, China. BALB/c nude mice (female, 6 weeks) were obtained from Vital River Laboratory Animal Technology Co., Ltd. (Beijing, China). Mice were housed in isolation and ventilation cages under SPF (Specific Pathogen Free) conditions in a climate-controlled room (25 ± 1.5 °C), 50 ± 10% humidity with 12/12 h light (7:30-19:30) and dark cycle, given unrestricted access to standard diet and water. To construct xenograft tumor models, 200 μL serum-free DMEM containing 1×10^7^ FaDu or Cal-27 cells were subcutaneously injected into the right flank of each mouse. Tumor volume was calculated using the following formula: V (volume) = (length × width^2^)/2. After mice were sacrificed, tumors were dissected, weighed, and resected for IHC staining.

#### Transfection, drug treatment and cell lysis

Transfection was carried out using Lipofectamine™ 2000 Transfection Reagent (Thermo Fisher Scientific) according to the manufacturer’s instruction. For inhibitors treatment, a DMSO stock solution of 200 mM phloretin (Sigma-Aldrich), 2 mM Tg101348 (Selleck) and 2 mM PD0325901 (Beyotime Biotechnology) was prepared. Before Vc or other small molecular treatment, cells were pretreated with 200 μM phloretin, 2 μM Tg101348 or 2 μM PD0325901 for 4 h. For Vc treatment, an aqueous stock solution of 50 mg/mL Vc (L-Ascorbic acid, Sigma-Aldrich) was prepared. Cells were incubated with or without Vc in culture medium for specific times and lysed immediately by cold RIPA (Solarbio) supplemented with 0.1% Protease Inhibitor Cocktail (Sigma-Aldrich), 1% Phosphatase Inhibitor Cocktail 3 (Sigma-Aldrich) and 0.5% Phosphatase Inhibitor Cocktail 2 (Sigma-Aldrich). The culture medium should not be renewed before Vc treatment for at least 12 h (for the newly addition of FBS will activate ERK rapidly and affect the accuracy of results).

#### Plasmid construction

m*Jak2* with Clover/mRuby2 tag and full-length coding sequences of mouse *Svct1* and *Svct2* were provided by Zhuo Han(Han et al., 2021). Both *Svct1* and *Svct2* cDNAs were inserted into pcDNA3.1(+) as follows: Restriction enzyme cutting site - kozak sequence - Flag tag-insertion - double Flag tag - Restriction enzyme cutting site.

m*Svct2* was either subcloned into pEGFPN1, pcDNA3-Clover or pcDNA3-mRuby2 using *pEASY*^®^-Basic Seamless Cloning and Assembly Kit (TransGen Biotech). All m*Svct2* mutants were produced by Site-directed mutation through splicing by overlap extension PCR and subcloned into pEGFPN1. The full-length coding sequences of mouse *Grb2* and *Ptpn12* were amplified from E14 mouse whole-brain cDNA, both were subcloned into pGFPC1, p3×Flag-CMV10, pCMV-HA, pcDNA3-Clover or pcDNA3-mRuby2 using *pEASY*^®^-Basic Seamless Cloning and Assembly Kit (TransGen Biotech). Mutants were produced by Site-directed mutation through splicing by overlap extension PCR. All inserted sequences were confirmed by sequencing (Sangon Biotech) and aligned with the NCBI database.

#### RNA interference and quantitative Real-Time PCR (qRT-PCR)

The siRNA sequences of m*Svct2* and m*Svct1* were derived from the study of Han et al.(Han et al., 2021). The siRNA sequence of h*Svct2* was exhibited in Supplementary materials. Cells were transfected with siRNA for 36 h, then the knockdown efficiency was determined by qRT-PCR and Western Blot. RNA was extracted with RNAiso Plus (Takara) following the manufacturer’s instruction. cDNA synthesized from 1 μg total RNA using PrimeScript™ RT reagent Kit with gDNA Eraser (Takara). Real-time PCR was performed using TB Green^®^ Premix Ex Taq™ II (Tli RNaseH Plus) (Takara). Values of gene expression were normalized to GAPDH expression using the ΔΔCT method.

#### Survival analysis and gene differentially expression analysis

Survival analysis and differentially expression of *SVCT1/2* in various cancers were performed using GEPIA2 web server (http://gepia2.cancer-pku.cn/#index), and the expression and clinical features were from the TCGA and the GTEx projects. *SVCT1/2* mRNA expression was classified as “Low Group” and “High Group” with preestablished cutoffs. The parameters of Kaplan-Meier survival plot and expression box plot were default. P-value less than 0.05 was considered statistically significant.

#### Immunofluorescence assay

Cells cultured on glass coverslips were fixed with 4% (w/v) paraformaldehyde (PFA) for 15 min and washed with PBS twice, permeabilized with 0.2% Triton X-100 in PBS for 15 min and washed with PBS, blocked with 1% FBS in PBS for 1 hour at room temperature. Then, cells were incubated with primary antibody at 4 ℃ overnight, washed with PBS, incubated with secondary antibody for 2 h at room temperature, and washed with PBS. Finally, mounted coverslips and photographed using FLUOVIEW FV3000 Confocal Laser Scanning Microscope (Olympus) with sequencial scan mode. Following antibodies were used: anti-Flag (1:500, Cat#F1804, Sigma-Aldrich), Goat anti-Mouse IgG (H+L) Highly Cross-Adsorbed Secondary Antibody, Alexa Fluor 555 (1:500, Cat#A-21424, ThermoFisher), Goat anti-Mouse IgG (H+L) Highly Cross-Adsorbed Secondary Antibody, Alexa Fluor 488 (1:500, Cat#A-11029, ThermoFisher).

#### Immunoprecipitation (IP) and Co-Immunoprecipitation (Co-IP)

Cells were transfected with specific plasmids for 36 h. Then cells were lysed using IP Lysis Buffer (25 mM Tris-HCl pH 7.4, 150 mM NaCl, 1% NP-40, 1 mM EDTA, 5% glycerol) supplemented with 0.1% Protease Inhibitor Cocktail (Sigma-Aldrich), 0.5% Phosphatase Inhibitor Cocktail 3 (Sigma-Aldrich) and 0.2% Phosphatase Inhibitor Cocktail 2 (Sigma-Aldrich) after washing with PBS twice (for samples treated with drugs for a very short time, the entire washing process should be controlled within 1 min). Each well of 6-well plate were lysed in 300 μL IP Lysis Buffer and incubate on ice for 20 min with periodic mixing. Cell lysate was centrifuged at 15000 × g for 10 min to pellet the cell debris at 4°C. Part of supernatant (40 μL) was transferred into a new tube used as positive control (input), the rest of supernatant (250 μL, IP) was transferred into another tube and incubated with anti-Flag antibody (1:100, Sigma-Aldrich) or control immunoglobulin G (IgG, Beyotime) at 4 °C for 6 h, with rotated mixing constantly. Then Flag tag was captured with Pierce Protein A/G Agarose (20 μL per sample, ThermoFisher) at room temperature for 2 h with mixing constantly. Supernatant was removed by centrifuging and washed with cold IP Lysis Buffer for 3 times, then bound proteins were eluted in 5 × SDS loading buffer by boiling for 10 min and analyzed by western blot. For muti-transmembrane protein like SVCT2, 4% CHAPS (Sigma-Aldrich) was added additionally in IP Lysis Buffer.

#### Cell apoptosis

Cell apoptosis was detected using Dead Cell Apoptosis Kit with Annexin V Alexa Fluor™ 488 & Propidium Iodide (PI) (Thermo Fisher Scientific) following manufacturer’s instructions. Unstained sample and single-stained sample were used as negative control or compensation. For B16-F10 cells and HeLa cells, BD FACSAria III flow cytometry system (Becton, Dickinson and Company) was used to signal collection and data was processed by FlowJo (Version 10.0.7) software. For FaDu cells and Cal-27 cells, ACEA NovoCyte 3130 flow cytometry system (ACEA Biosciences Inc.) was used to signal collection and data processing.

#### Western Blot

Proteins were separated by electrophoresis on 10% Sodium Dodecyl Sulfate Poly-Acrylamide Gel (SDS-PAGE) and transferred to polyvinylidene difluoride (PVDF) membranes (0.22 μm). The details of Western Blot were performed following Protocols and Troubleshooting on Advansta’s website. The membranes were detected using WesternBright ECL kit (Advansta) or WesternBright Sirius Chemiluminescent Detection Kit (Advansta) on ChemiDoc™ MP gel imaging analysis system (BioRad).

The primary antibodies and dilutions were used as follows:

Anti-Flag (1:3000, Cat#F1804, Sigma-Aldrich), anti-GFP (1:1000, Cat#66002-1-Ig, Proteintech), anti-HA (1:1000, Cat#PA1-985, ThermoFisher), anti-JAK2 (1:250, Cat#sc-390539, Santa Cruz), anti-pJAK2 (1:500, Cat#3771, Cell Signaling Technology), anti-ERK1/2 (1:3000, Cat#4695, Cell Signaling Technology), anti-pERK1/2 (1:5000, Cat#9101, Cell Signaling Technology), anti-PTPN12 (1:500, Cat#sc-271351, Santa Cruz), anti-GAPDH (1:3000, Cat#HC301, TransGen), anti-β-Tubulin (1:3000, Cat#HC101, TransGen), anti-SVCT2 (1:1000, Cat#ab229802, Abcam), anti-phosphotyrosine (1:800, Cat#9411, Cell Signaling Technology), anti-phosphoserine (1:250, Cat#sc-81514, Santa Cruz).

The secondary antibodies were used as follows:

HRP-labeled Goat Anti-Rabbit IgG(H+L) (A0208, Beyotime Biotechnology), HRP-labeled Goat Anti-Mouse IgG(H+L) (A0216, Beyotime Biotechnology).

#### Dual-Luciferase Reporter (DLR) Assay

Dual-Luciferase Reporter assays were performed with a dual luciferase reporter assay system (TransGen Biotech) following the manufacturers instructions. Briefly, the pathway reporter plasmid pSRE -luc and *Renilla* luciferase control plasmid pRL-TK were co-transfected into F9 cells or HEK293T cells. 36 h after transfection, cells were lysed with 250 μl/well (12-well plate) cell lysis buffer for 10 min with shaking. Then centrifuged at 15000 × g for 10 min to pellet the cell debris at 4°C. 20 μl of each sample was transferred into a 96-well plate (Corning) and assayed by *TransDetect*^®^ Double-Luciferase Reporter Assay Kit. Data were collected by Varioskan LUX (ThermoFisher).

#### Measurement of intracellular Vc content

Measurement of intracellular Vc content by HPLC was totally performed following the instructions in previous reports(Han et al., 2021).

#### Microscale Thermophoresis (MST)

MST is an optical method to characterize protein and small-molecule interactions in biological liquids. All MST experiments are setup with one fluorescently labeled protein (Target, SVCT2 and SVCT2 mutants) at a fixed concentration that is mixed with various concentrations of a non-fluorescent molecule (Ligand, Vc) according to instructions provided by Nanotemper.

SVCT2-GFP and other mutants were transfected into HEK293T cells for 36 h. Cells (60 mm culture dish) were washed with PBS twice and extracted in 700 μL MST buffer (25 mM Tris-HCl pH 7.4, 150 mM NaCl, 1% NP-40, 1 mM EDTA, 5% glycerol and 0.05% Tween-20) supplement with protease inhibitors. Cell debris was removed by centrifugation at 15000 × g for 10 min at 4°C.

L-ascorbic acid sodium (Solarbio) stock solution (2 M) was diluted in MST buffer. The serial dilution steps follow the instructions in MO.Affinity Analysis v2.3. The ligand concentration is from 1000 mM to 0.0305 mM and bubbles should be avoided while mixing target and ligand. Then binding affinity between Vc and SVCT2s was detected by monitoring binding induced MST signal changes. All interactions were measured in Monolith NT.115 Standard Treated Capillaries (MO-K002, Nanotemper) at 25℃ with an MST-on time of 20 seconds and an MST-off time of 5 seconds. Measurements were performed using a NanoTemper^®^ Technologies Monolith^®^ NT.115 system and data was analyzed using MO.Affinity Analysis v2.3.

#### Förster Resonance Energy Transfer (FRET)

The FRET efficiency between proteins were measured using sensitized emission FRET, mRuby2 being used as a donor and Clover as an acceptor(Lam et al., 2012). Each DNA sequence was inserted into either pCDNA3-mRuby2 (Addgene#40260) or pCDNA3-Clover (Addgene#40259). To establish FRET assay, HEK293T cells were transfected with indicated plasmids for 36 h, then cells were cultured into 35 mm cell culture dish with glass bottom (NEST) at the desired density. When reach 60-80% confluency, cells were photographed using FLUOVIEW FV3000 Confocal Laser Scanning Microscope (Olympus), and sensitized emission FRET analysis was performed using cellSens Dimension V3.1 (Olympus).

Image acquisition: For the acquisition of fluorescence images for FRET, three different combinations of excitation and emission parameters was set in the confocal microscope: FRET channel (the sample is illuminated with the donors excitation light 488 nm and the accepto rs emission light 560 -630 nm is detected), Donor channel (the sample is illuminated with the donors excitation light 488 nm and the donors emission light 510-550 nm is detected) and Acceptor channel (the sample is illuminated with the acceptors excitat ion light 561 nm and the acceptors emission light 560 -630 nm is detected). Meanwhile, the sequential scan option was “None”. Then, two images of a sample that only contains the donor (FRET channel and Donor channel) and two images of a different sample that only contains the acceptor (FRET channel and Acceptor channel) were snapped for FRET correction. The parameters for FRET correction images were same with the acquisition of images for FRET.

FRET correction: The appropriate reference images were loaded to cellSens Dimension V3.1 and FRET correction was performed using *FRET Correction* dialog box. After selected the appropriate reference images, background and at least 5 different ROIs, and FRET correction is calculated and correction factors DSBT a (FRET channel/Donor channel) and ASBT b (FRET channel/Acceptor channel) were recorded. Finally, correction factors are the mean of 3 replicates.

FRET analysis: The FRET images were loaded to cellSens Dimension V3.1 and FRET analysis was performed using *FRET analysis* dialog box. Then appropriate channels were selected and a ROI as background was defined. The computation method was *Xia* in the *method* list and this computation method is taken from previous publication(Xia and Liu, 2001). Next, correction factors were loaded and FRET analysis was performed. According to cellSens manual, the FRET map resulting from the Xia method is a gray-value image which automatically has a predefined pseudo color table. High values are displayed in red with this pseudo color table and low ratio values are displayed in magenta.

#### Time-Resolved FRET (TR-FRET)

Time-Resolved FRET is based on sensitized emission FRET. HEK293T cells were transfected with indicated plasmids for 36 h, then cells were cultured into 35 mm cell culture dish with glass bottom (NEST) and cultured for 10 h. Before FRET images were snapped, half of culture medium was removed into a new centrifuge tube, and Vc stock solution was added up to 2-fold indicated concentrations. Cell culture dish was placed under the confocal microscope and FRET images were snapped with time stack. The first three images were collected without Vc treatment, in the interval between the third image and the fourth image, the culture medium premixed with Vc was added back to culture dish carefully. Quantitation was done using *Intensity profiles* command in cellSens Dimension V3.1 (Olympus).

#### Silver stain assay

Proteins were separated by 10% SDS-PAGE. The silver stain assays were performed using PAGE Gel Silver Staining Kit (Solarbio) following the manufacturer’s instructions. But after each step, the gel was rinsed with deionized water.

#### Transwell assay

1×10^5^ cells in DMEM were seeded into a Transwell chamber (NEST) which loaded with 8 μm small-pore polycarbonate membrane, the cell-free lower chamber was added with DMEM containing 10% FBS. After the migration, cells on Transwell chamber were fixed with PBS containing 4% PFA for 30 min at room temperature, and washed twice with PBS. The polycarbonate membranes were wiped gently with a fluffy cotton slightly to swab cells in the upper layer, permeabilized with methanol for 5 min, washed once with PBS, and stained with crystal violet (Sangon Biotech) staining solution (10 times dilution of 1% DMSO crystal violet stock solution by PBS) at 37°C for 30 min. Finally, the chambers were rinsed with PBS 3 times to remove excess dye, and photographed by inverted microscope (Nikon).

#### Phosphoproteome

Quantitative phosphoproteome of Vc-stimulated F9 cells was performed using multiplex tandem mass tag (TMT)-labeling coupled with liquid chromatography-mass spectrometry (LC-MS). Briefly, F9 cells were treated with Vc (50 μg/mL) for 20 min, then cells were lysed in lysis buffer containing 100 mM NH_4_HCO_3_ (pH=8),6 M Urea, 0.2% SDS and 1% Phosphatase Inhibitor Cocktail 3 (Sigma-Aldrich), followed by quick freezing in liquid nitrogen. Phosphoproteome was performed by Novogene (Beijing, China) and the .raw data was searched using Mus_musculus_uniprot_2019.01.18.fasta (85165 sequences) database by Proteome Discoverer 2.2 (ThermoFisher).

#### Phosphatase assay

HEK293T cells were transfected with Flag-tagged PTPN12 and PTPN12 mutants, proteins were immunoprecipitated using IP lysis buffer supplement with 1 mM PMSF (Solarbio) and anti-Flag antibody (Sigma-Aldrich) to verify the equal amounts of immunoprecipitated protein. Then precipitates were analyzed using pNPP as a substrate as described(Streit et al., 2006).

#### Intratumoral injection (i.t.) and PD treatment

Tumor-bearing mice were randomly assigned to 5 groups: Control, Vc (Low Dose), Vc (High Dose), DMSO, PD. Intratumoral injection of L-ascorbic acid sodium (Solarbio) was administered when the longer diameters of tumors reached 5 millimeters. For B16-F10 xenograft tumor model, mice from Vc (High Dose) group were given at doses of 100 mg/mL L-ascorbic acid sodium aqueous solution followed by 50 μL/0.5 cm^3^ tumor volume every 2 days, mice from Vc (Low Dose) group were given at doses of 2.5 mg/mL L-ascorbic acid sodium followed by 50 μL/0.5 cm^3^ tumor volume every 2 days and mice from Control group were given sodium chloride solution every 2 days. For FaDu/Cal-27 xenograft tumor model, mice from Vc (High Dose) group were given at doses of 200 mg/mL L-ascorbic acid sodium aqueous solution followed by 50 μL/0.5 cm^3^ tumor volume every 2 days, mice from Vc (Low Dose) group were given at doses of 12.5 mg/mL L-ascorbic acid sodium followed by 50 μL/0.5 cm^3^ tumor volume every 2 days and mice from Control group were given sodium chloride solution every 2 days. To balance the osmotic effect of high-dose Vc, all mice from Control and Vc (Low Dose) group were given the same molar concentration of sodium with Vc (High Dose) group. Intragastric administration of DMSO (Sigma-Aldrich) or PD0325901 (Beyotime Biotechnology) was administered from day 3 after inoculation, 10 mg/kg, alternating days. Mice were sacrificed until the tumor size humane endpoint was met.

#### Histology and IHC

Tumors were fixed in 4% PFA, dehydrated, paraffin embedded and sliced (8 μm). After being baked at 37℃, they were placed at room temperature. Slices were deparaffinized in xylene (2 times, 5 min each), and rehydrated with successive washes in 100%, 96%, 80%, and 70% ethanol. The Slices were stained with H&E for quality control. For IHC, it was performed following the standard procedures, and following antibodies were used: anti-pERK (Cat#9101, Cell Signaling Technology), anti-Ki-67 (Cat#RMA-0731, MXB Biotechnologies), anti-E-Cadherin (Cat#20874-1-AP, proteintech), anti-N-Cadherin (Cat#13116, Cell Signaling Technology), anti-Vimentin (Cat#10366-1-AP, proteintech). Slices were scanned using Pannoramic SCAN 150 (3DHistech) and processed using CaseViewer v2.2 (3DHistech).

#### Quantification and statistical analysis

Statistical details of each experiment can be found in the Figure legends. The number of experimental replicates is indicated in the Figure legends or outlined in the Method details section. Statistical comparisons were two-tailed Student’s *t*-test as described in the figure legends using GraphPad Prism software (version 6.07) or Excel. Data presented in graphs are mean ± SD from at least three replicates unless stated otherwise. Statistical significance was set at ∗ indicates p<0.05, ∗∗ indicates p<0.01, ∗∗∗ indicates p<0.001.

