## Supplementary material for "Vitamin C functions as double-edge sword on cancer progression depending on ERK activation or inhibition mediated by its receptor SVCT2": Figure S1-S9 and Table S1-S3

### SUPPLEMENTARY DATA

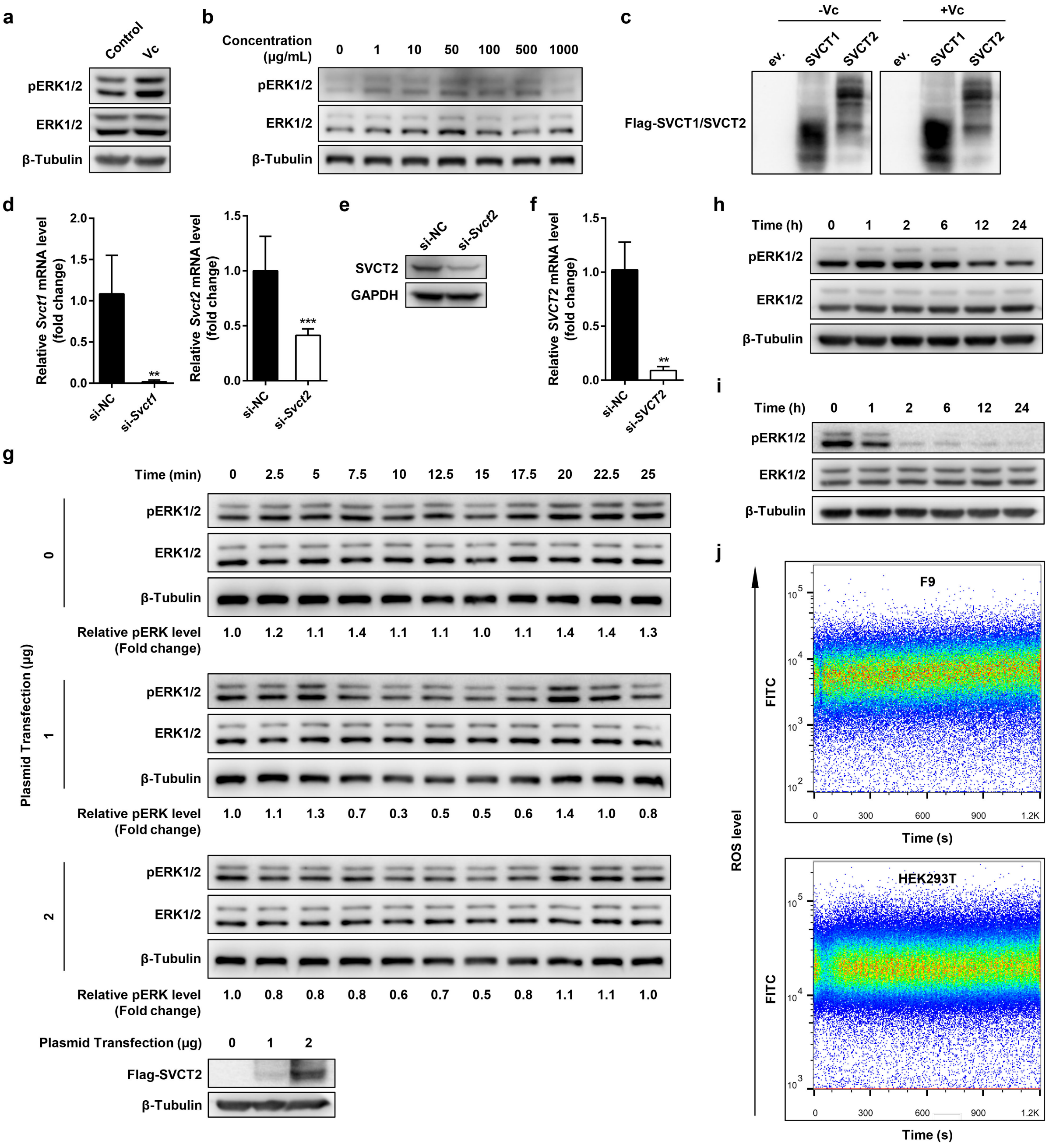


**Figure S1. Vc induces ERK activation in an SVCT2-dependent manner** (related to Figure 1)

**(a)** RAW264.7 macrophages were treated with Vc (50 μg/mL) and lysed at 20 min. Lysates were immunoblotted for endogenous pERK1/2, total ERK1/2 and β-Tubulin.

**(b)** Jurkat T lymphocyte leukemia cells were treated with Vc of various concentrations as indicated and lysed at 20 min. Lysates were immunoblotted for endogenous pERK1/2, total ERK1/2 and β-Tubulin.

**(c)** Validation of plasmids expressions. F9 cells were transfected as indicated for 36 h and treated with Vc (50 μg/mL) for 20 min. Lysates were immunoblotted for Flag-tagged SVCT1 or Flag-tagged. e.v., empty vector.

**(d and e)** F9 cells were transfected with *Svct1*-siRNA or *Svct2*-siRNA, cell lysates were harvested 36 h later, decreased levels of endogenous *Svct1* and *Svct2* mRNA was verified by qPCR (**d**) and decreased expression of endogenous SVCT2 was verified by immunoblots (**e**).

**(f)** HEK293T cells were transfected with *SVCT2*-siRNA, cell lysates were harvested 36 h later, and decreased level of endogenous *SVCT2* mRNA was verified by qPCR.

**(g)** HEK293T cells were transfected with different doses of Flag-SVCT2 plasmid as indicated and subcultured. 36 h after transfection, cells were treated with Vc (50 μg/mL) and lysed at indicated times. Lysates were immunoblotted for endogenous pERK1/2, total ERK1/2, β-Tubulin and Flag-tagged SVCT2.

**(h and i)** F9 cells (**h**) and HEK293T cells (**i**) were treated with Vc (1000 μg/mL) and lysed at indicated times. Lysates were immunoblotted as indicated.

**(j)** Reactive oxygen species (ROS) levels detected by flow cytometry. F9 cells or HEK293T cells were treated with Vc (500 μg/mL) and intracellular ROS levels were detected immediately in real-time. The detection lasted for 20 min.

Error bars represent SD of three independent experiments. **, $P<0.01$; ***, $P<0.001$; two-tailed Student's *t*-test.

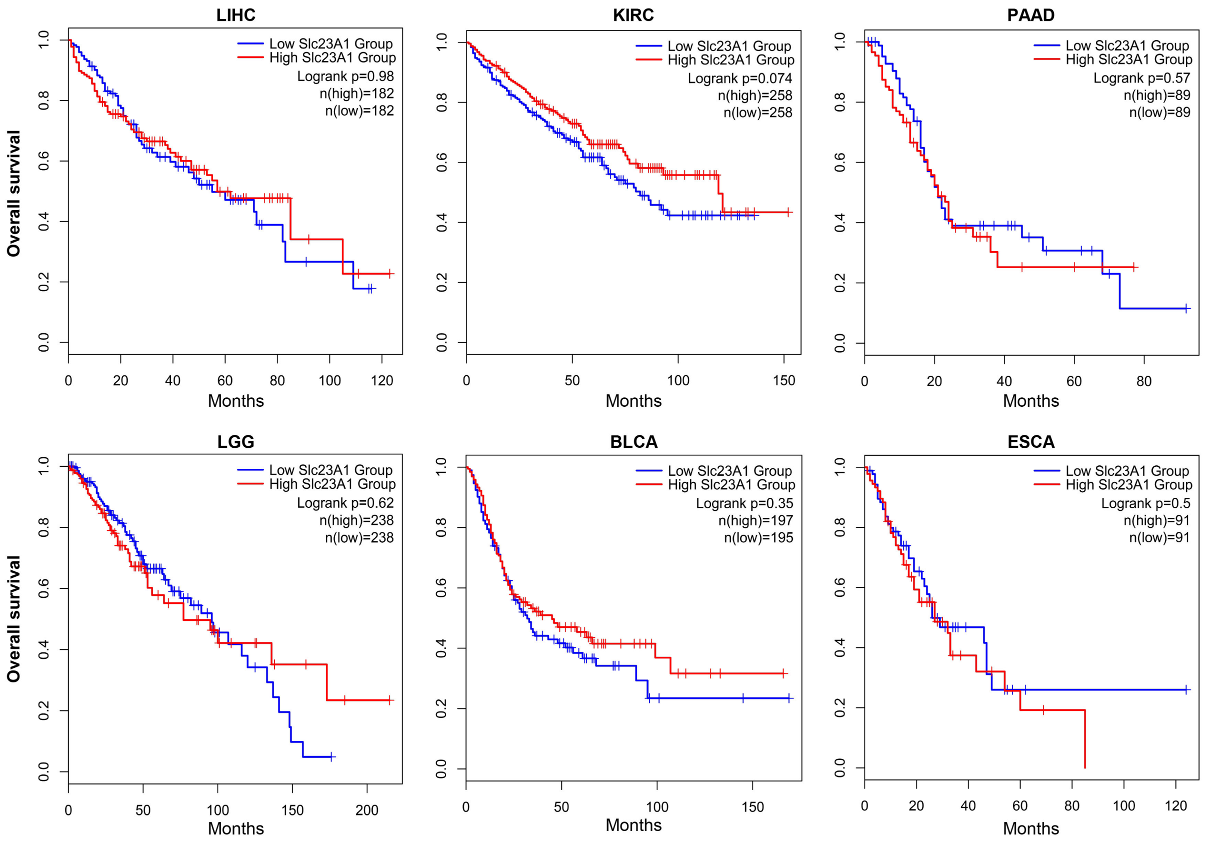


**Figure S2. Survival and differentially expression analysis of SVCT1 (SLC23A1) in various cancers** (related to Figure 2)

Differential expression and survival analysis of SVCT1 (SLC23A1) in various cancers were performed using GEPIA2 web server (http://gepia2.cancer-pku.cn/#index), and the expression and clinical features were from the TCGA and the GTEx projects.

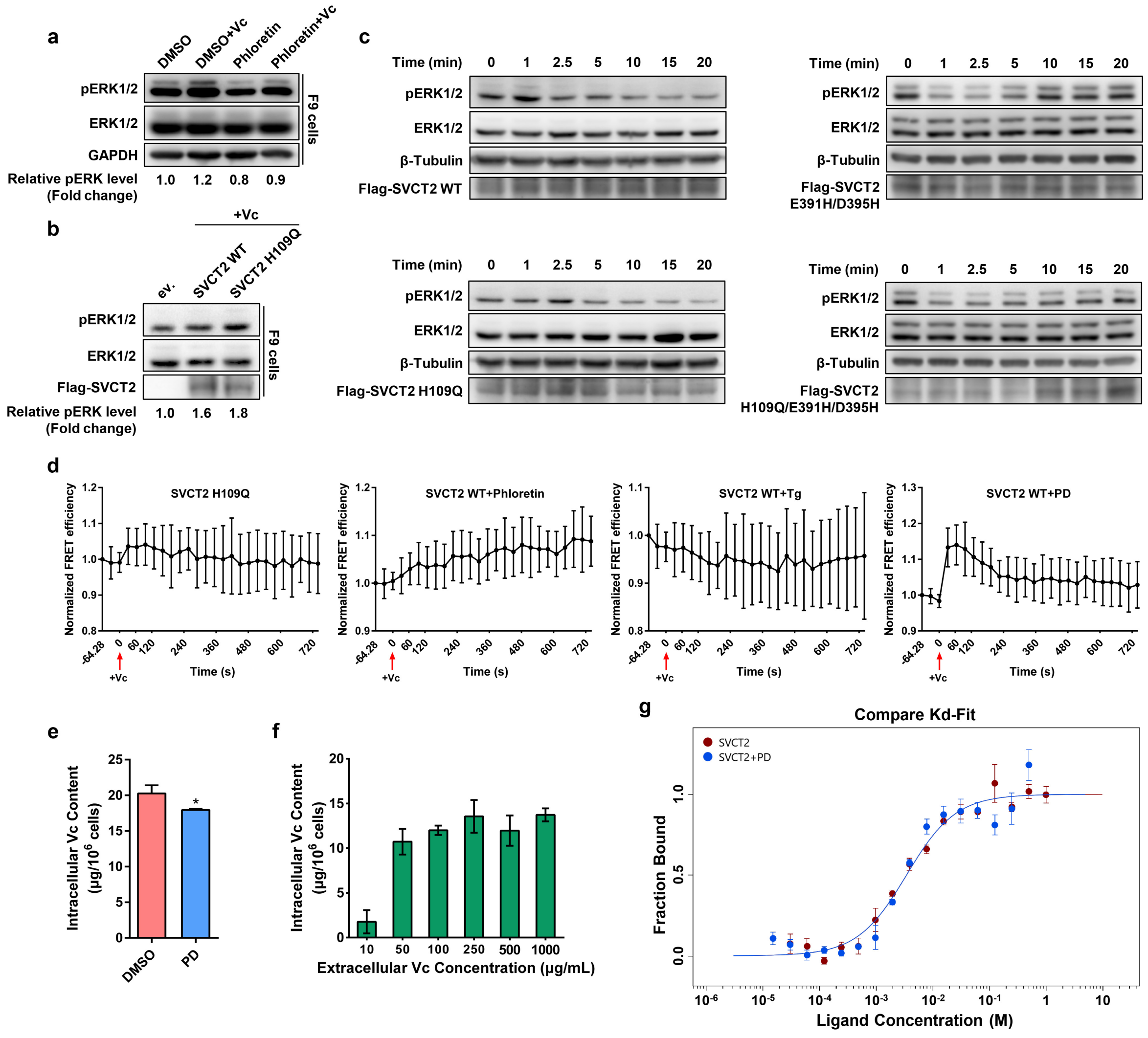


**Figure S3. Extracellular Vc binding to SVCT2 is sufficient for Vc-mediated ERK activation** (related to Figure 3)

**(a)** F9 cells were pretreated with phloretin (200 μM) or DMSO for 4 h, then cells were treated with Vc (50 μg/mL) for 20 min, and lysates were immunoblotted as indicated.

**(b)** F9 cells were transfected as indicated for 36 h and then treated with Vc (50 μg/mL) for 20 min. Lysates were immunoblotted as indicated. ev., empty vector.

**(c)** HEK293T cells were transfected with SVCT2 WT, SVCT2 H109Q, SVCT2 E391H/D395H or SVCT2 H109Q/E391H/D395H for 36 h, and then treated with Vc (500 μg/mL) for the indicated times. Lysates were immunoblotted as indicated.

**(d)** Time-Resolved FRET in living cells. HEK293T cells were co-transfected with SVCT2-Clover and SVCT2-mRuby2 or SVCT2 H109Q-Clover and SVCT2 H109Q-mRuby2 as indicated for 36 h, then cells were treated with Phloretin (200 μM), Tg (2 μM) or PD (2 μM) respectively for 4 h and photographed time-scale images using confocal microscope for sensitized emission FRET analysis. For one time-scale, cells were treated with Vc (500 μg/mL) immediately after 3 pictures were snapped. All data were mean ± SD of the relative FRET efficiency from at least 9 cells (SVCT2 H109Q, n=10; SVCT2 WT+Phloretin, n=9; SVCT2 WT+Tg, n=10; SVCT2 WT+PD, n=14). In SVCT2 WT+PD, the maximum response after Vc treatment was significantly different to that of before Vc treatment ($P<0.01$, two-tailed Student's *t*-test).

**(e)** Determination of intracellular Vc content by HPLC. F9 cells were pretreated with 2 μM PD for 4 h. Then cells were exposed to Vc with initial concentration of 50 μg/mL and incubated for 6 h. Intracellular Vc content was determined by HPLC.

**(f)** Determination of intracellular Vc content by HPLC. F9 cells were treated with various concentrations of Vc as indicated for 6 h. Then intracellular Vc content was determined by HPLC.

**(g)** Dose-response curve for the binding interaction between SVCT2 and Vc with or without PD treatment by MST analysis (Kd model). The concentration of GFP-fused SVCT2/SVCT2 mutants in HEK293T lysate was constant, while the concentration of the L-ascorbate sodium between 1000 mM to 0.0305 mM. Ligand concentrations on the x-axis were plotted in mole (M). The dose-response curve of SVCT2-Vc without PD (red) shared the same data with Figure 3**d**.

Error bars represent SD of at least three independent experiments. *, $P<0.05$; two-tailed Student's *t*-test.

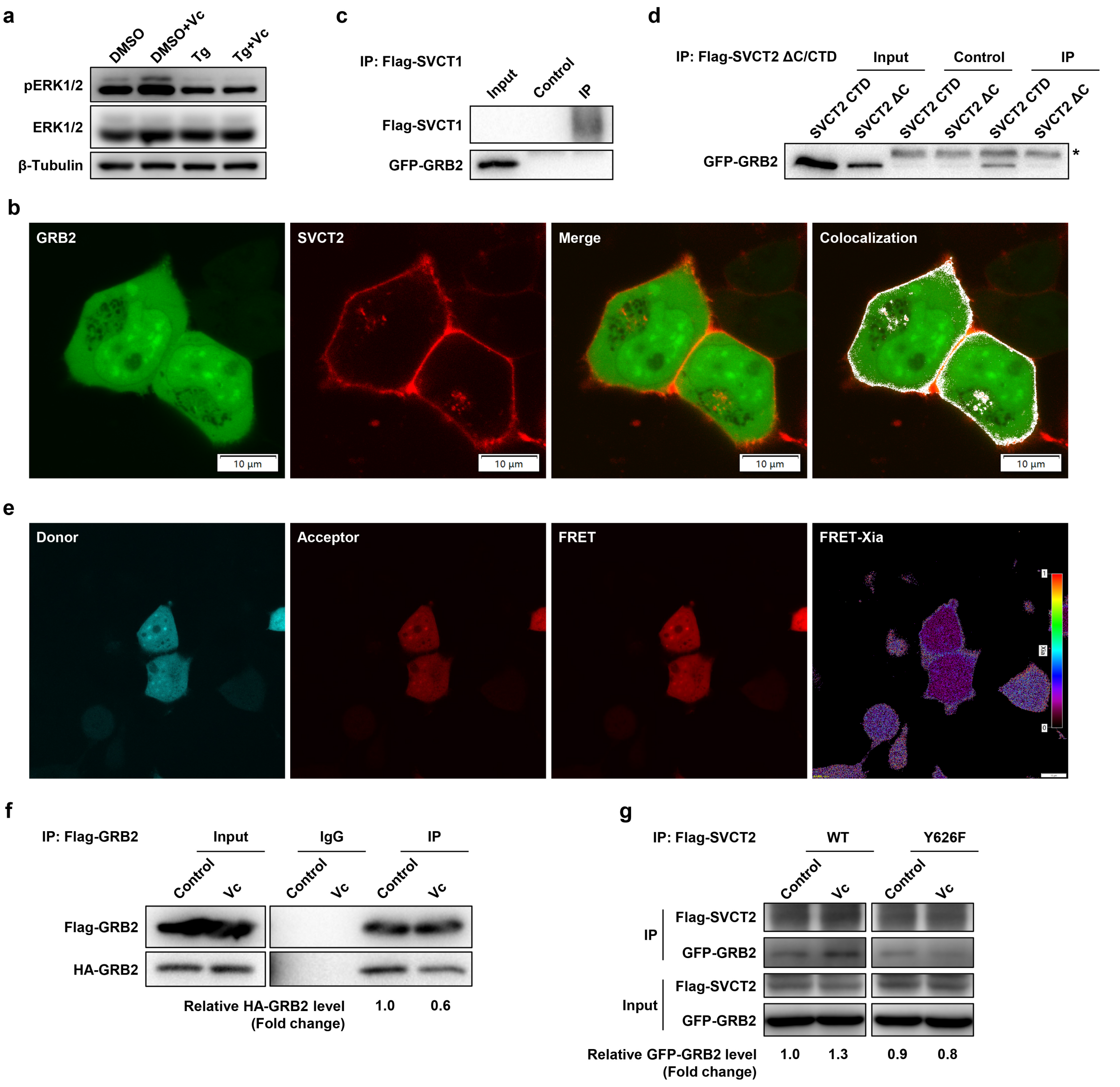


**Figure S4. The JAK2-GRB2 axis contributes to Vc-induced ERK activation** (related to Figure 4)

**(a)** HEK293T cells were pretreated with Tg (2 μM) or DMSO for 4 h, then cells were treated with Vc (50 μg/mL) for 20 min. Lysates were immunoblotted as indicated.

**(b)** The colocalization between GFP-GRB2 and Flag-tagged SVCT2 in HEK293T cells. Scale bar, 10 μm.

**(c and d)** HEK293T cells were co-transfected with Flag-tagged SVCT1 (**c**), SVCT2ΔC (**d**) or SVCT2 CTD (**d** and GFP-GRB2 respectively for 36 h. Flag-tag was immunoprecipitated and precipitates were blotted for GFP-tagged GRB2. *, heavy chain of anti-Flag antibody. Control, pCDNA3.1(+) fused with Flag tag were co-transfected with GFP-GRB2, Flag-tag was immunoprecipitated and precipitates were blotted for GFP-tagged GRB2.

**(e)** Representative confocal images of GRB2-Clover (donor), GRB2-mRuby2 (acceptor), FRET and FRET map. HEK293T cells were co-transfected with GRB2-Clover and GRB2-mRuby2 for 36 h, then cells were photographed using confocal microscope for sensitized emission FRET analysis. The FRET map resulting from the Xia method was displayed as pseudo color table. High values were displayed in red and low ratio values were displayed in magenta. Scale bar, 10 μm.

**(f)** HEK293T cells were co-transfected with Flag-tagged GRB2 and HA-tagged GRB2. 36 h after transfection, cells were treated with Vc (50 μg/mL) for 20 min. Flag-GRB2 was immunoprecipitated and precipitates were blotted for HA-tagged GRB2. IgG, immunoglobulin G.

**(g)** HEK293T cells were co-transfected with Flag-tagged SVCT2 WT/SVCT2 Y626F and GFP-GRB2 for 36 h, then cells were treated with Vc (50 μg/mL) for 2 h. Flag-SVCT2/SVCT2 Y626F was immunoprecipitated and precipitates were blotted for GFP-tagged GRB2.

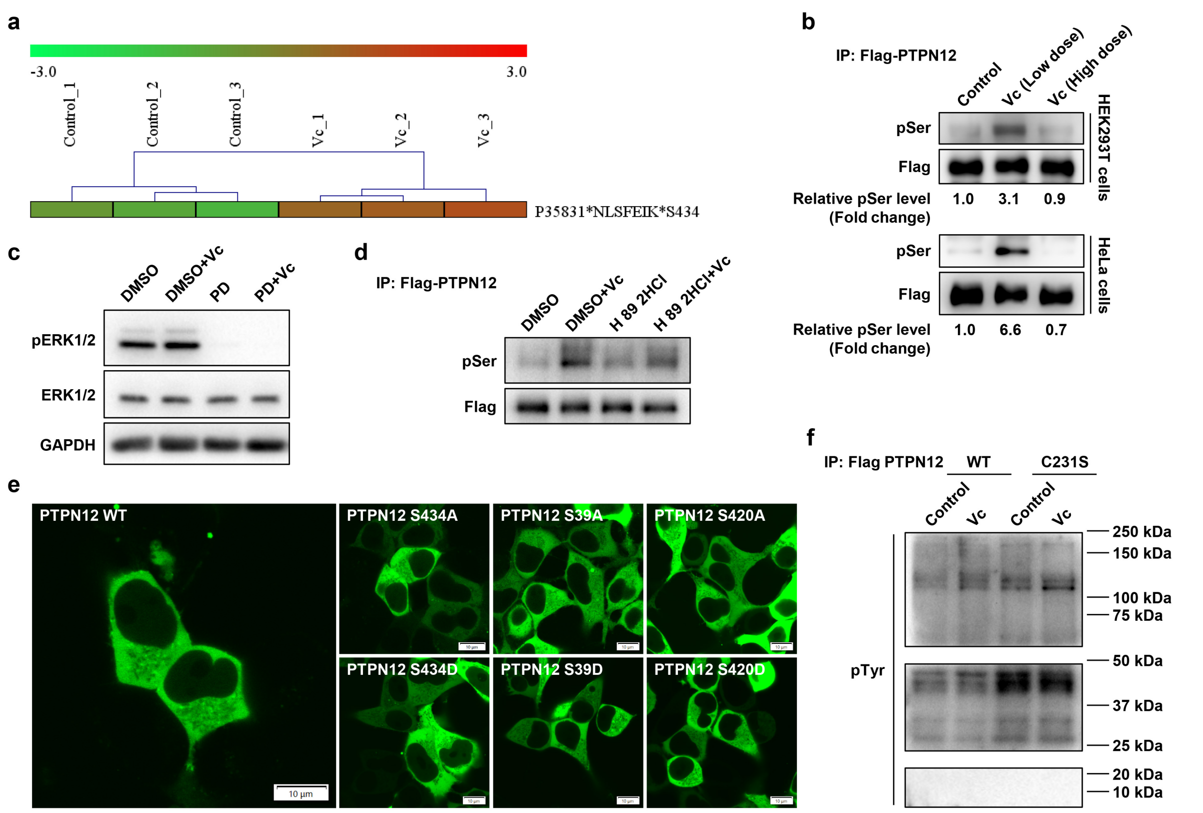


**Figure S5. Vc-induced ERK activation phosphorylates PTPN12 at Ser$^{434}$ to inhibit its phosphatase activity** (related to Figure 5)

**(a)** Quantitative phosphoproteome of Vc-stimulated F9 cells indicated the phosphorylation of PTPN12 at Ser$^{434}$ was induced by Vc.

**(b)** HEK293T or HeLa cells were transfected with Flag-tagged PTPN12 for 36 h, then cells were treated with low-dose (50 μg/mL) or high-dose (500 μg/mL) of Vc and lysed at indicated times. Flag-tagged PTPN12 was immunoprecipitated, and precipitates were blotted for phospho-serine. pSer, phospho-serine.

**(c)** F9 cells were pretreated with PD (2 μM) or DMSO for 4 h, and then cells were treated with Vc (50 μg/mL) for 30 min. Lysates were immunoblotted as indicated.

**(d)** HEK293T cells were transfected with Flag-tagged PTPN12. 36 h after transfection, cells were pretreated with H 89 2HCl (30 μM) (a specific inhibitor of PKA) or DMSO for 1 h. Then cells were treated with Vc (50 μg/mL) for 20 min. Flag-tagged PTPN12 was immunoprecipitated, and precipitates were blotted for phospho-serine. pSer, phospho-serine.

**(e)** Subcellular localization of PTPN12 mutants in HEK293T cells. HEK293T cells were transfected with Flag-PTPN12 and Flag-PTPN12 mutants for 36 h. Then the localization of PTPN12 mutants was examined by immunofluorescence assay. Cells were photographed under confocal microscope system. Scale bar, 10 μm.

**(f)** HEK293T cells were transfected with Flag-tagged wild-type (WT) PTPN12 or PTPN12 C231S for 36 h, cells were treated with Vc (50 μg/mL) for 20 min and lysed immediately. Flag-tagged PTPN12 was immunoprecipitated, and precipitates were blotted for phospho-tyrosine.

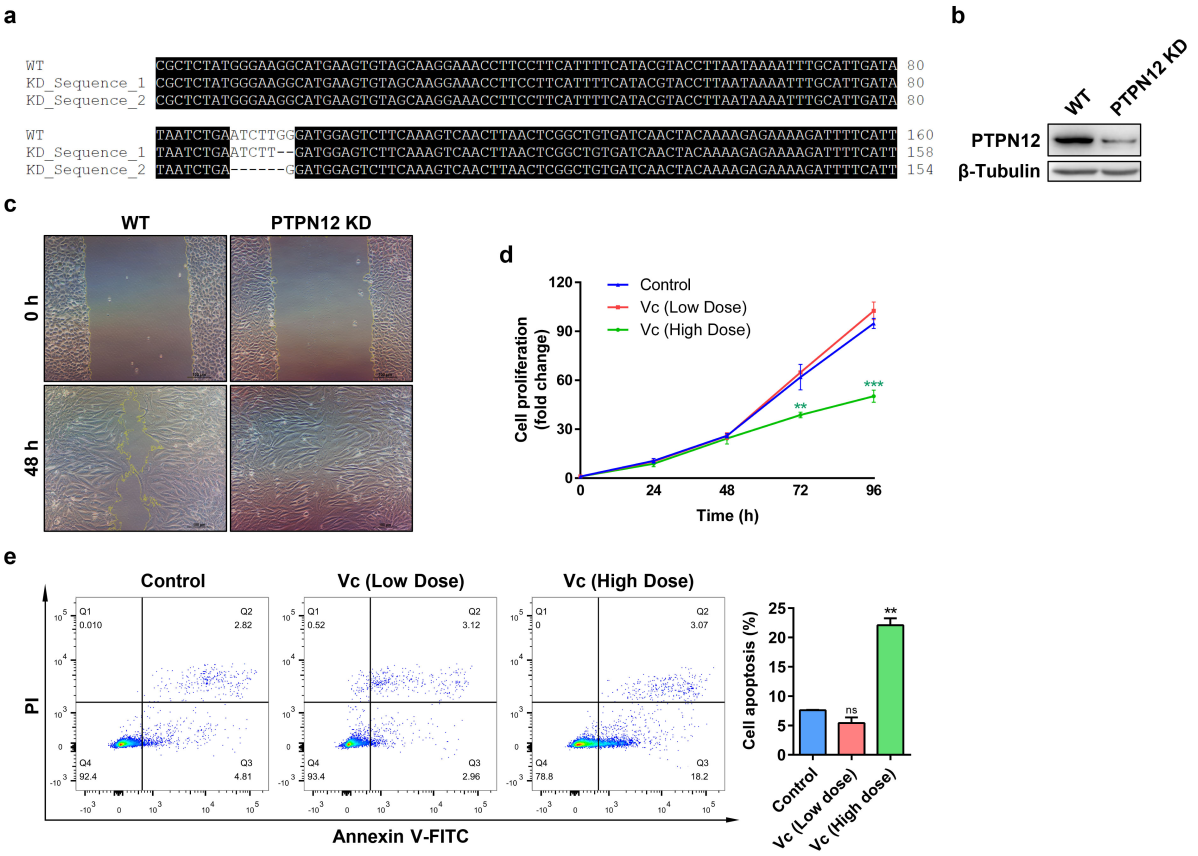


**Figure S6. *Ptpn12* knockdown affects the progression of cancer** (related to Figure 6)

**(a)** DNA sequencing and alignment of wild-type B16-F10 cells (WT) and *Ptpn12* knockdown B16-F10 cells (PTPN12 KD). The alignment was performed in BioXM (v2.6.0).

**(b)** Wild-type B16-F10 cells and *Ptpn12* knockdown B16-F10 cells were lysed and lysates were immunoblotted for PTPN12 and β-Tubulin.

**(c)** The migration ability of wild-type B16-F10 cells and *Ptpn12* knockdown B16-F10 cells were detected by wound healing assay. Scale bar, 100 μm.

**(d)** Proliferation of *Ptpn12* knockdown B16-F10 cells treated with different doses of Vc. Low dose, 10 μg/mL. High dose, 500 μg/mL.

**(e)** Apoptosis of *Ptpn12* knockdown B16-F10 cells treated with different doses of Vc were detected by flow cytometry. Results were representative of one of three similar experiments. Low dose, 10 μg/mL. High dose, 500 μg/mL.

For all graphs, data were presented as mean ± SD. ns, not significant; **, P<0.01; ***, P<0.001; two-tailed Student's *t*-test compared to control.

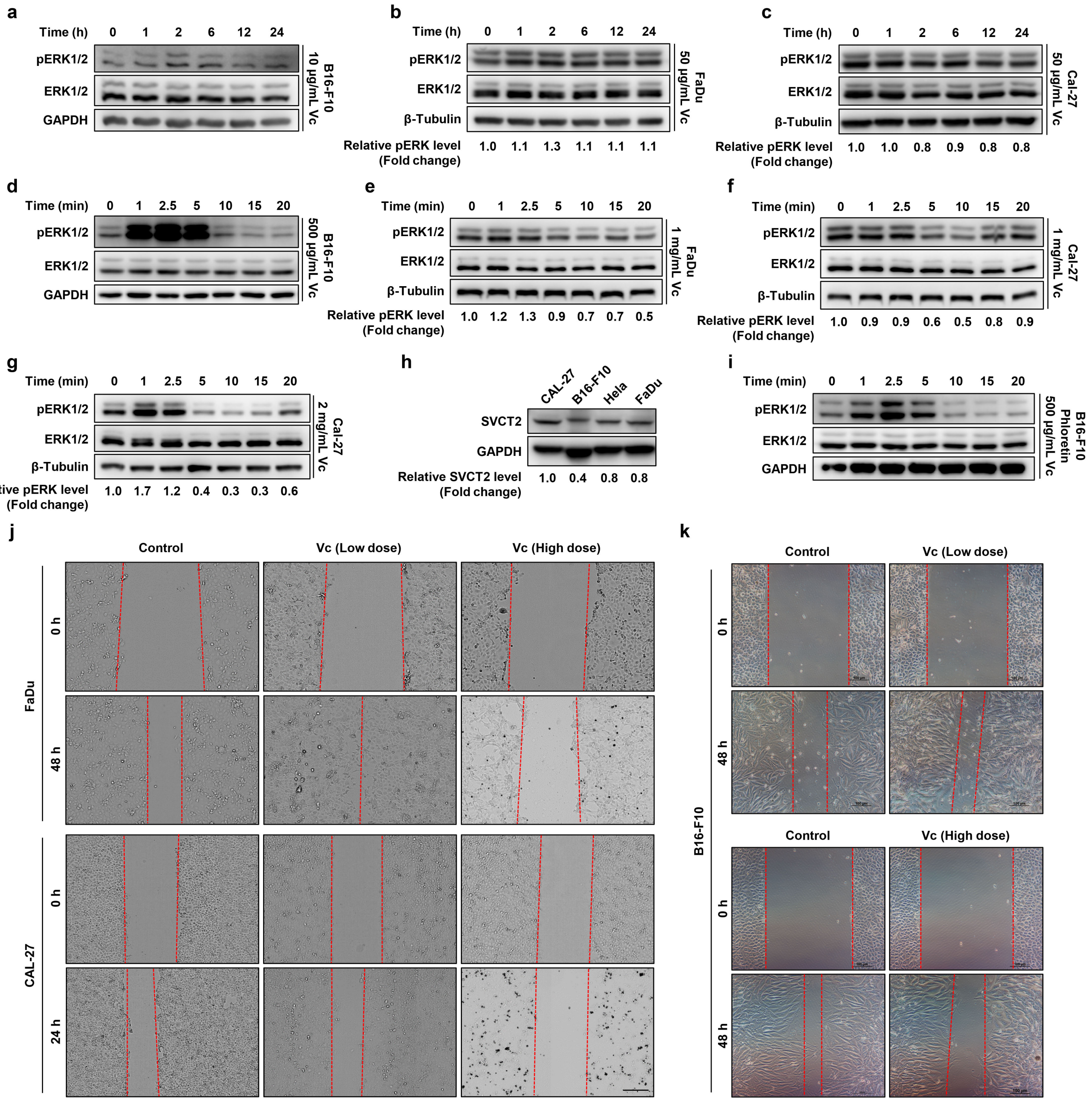


**Figure S7. High-dose Vc affects the phenotype of cancer cells through inhibiting ERK** (related to Figure 7)

**(a-g)** B16-F10, FaDu or Cal-27 cells were treated with Vc at indicated concentrations and lysed at indicated times. Lysates were immunoblotted as indicated.

**(h)** Cal-27, B16-F10, HeLa and FaDu cells were lysed using IP lysis buffer and lysates were immunoblotted for endogenous SVCT2 and GAPDH.

**(i)** B16-F10 cells were pretreated with phloretin (200 μM) for 4 h, and then treated with Vc (500 μg/mL) and lysed at indicated times. Lysates were immunoblotted as indicated.

**(j and k)** The migration ability of FaDu, Cal-27 and B16-F10 cells were detected by cell wounded assay. Scale bar, 100 μm.

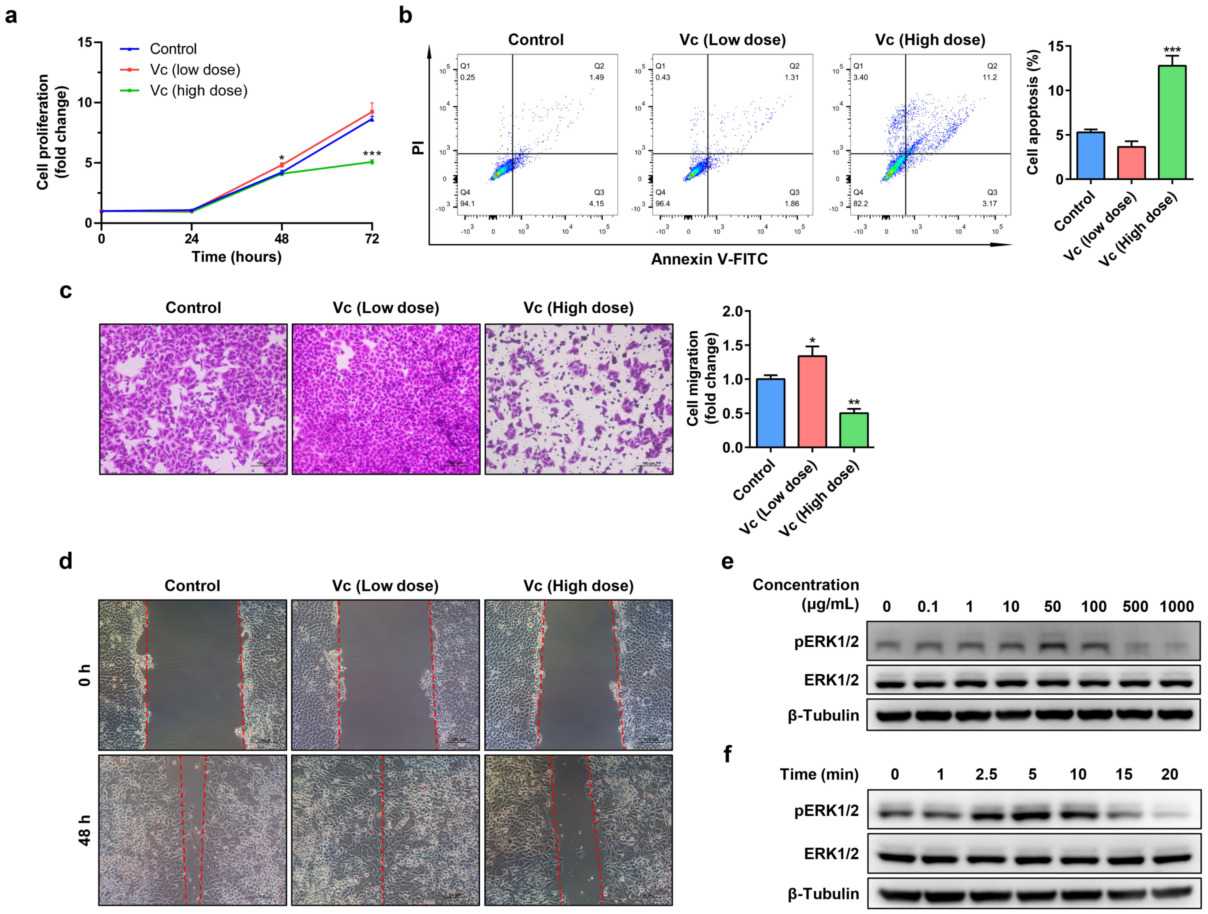


**Figure S8. Vc affects the phenotype of HeLa cells**

**(a)** Proliferation of HeLa cells treated with different doses of Vc were evaluated by cell counting.

**(b)** Apoptosis of HeLa cells treated with different doses of Vc were detected by flow cytometry.

**(c and d)** The migration ability of HeLa cells treated with different doses of Vc were detected by transwell assay (**c**) and wound healing assay (**d**).

**(e)** HeLa cells were treated with Vc at indicated concentrations (0-1000 μg/mL) for 20 min. Lysates were immunoblotted as indicated.

**(f)** HeLa cells were treated with Vc (1000 μg/mL) and lysed at indicated times. Lysates were immunoblotted as indicated.

In Figure S8, low dose Vc was 50 μg/mL; high dose was 1000 μg/mL. Error bars represent SD of three replicates. *, P<0.05; **, P<0.01; ***, P<0.001; two-tailed Student's *t*-test compared to control.

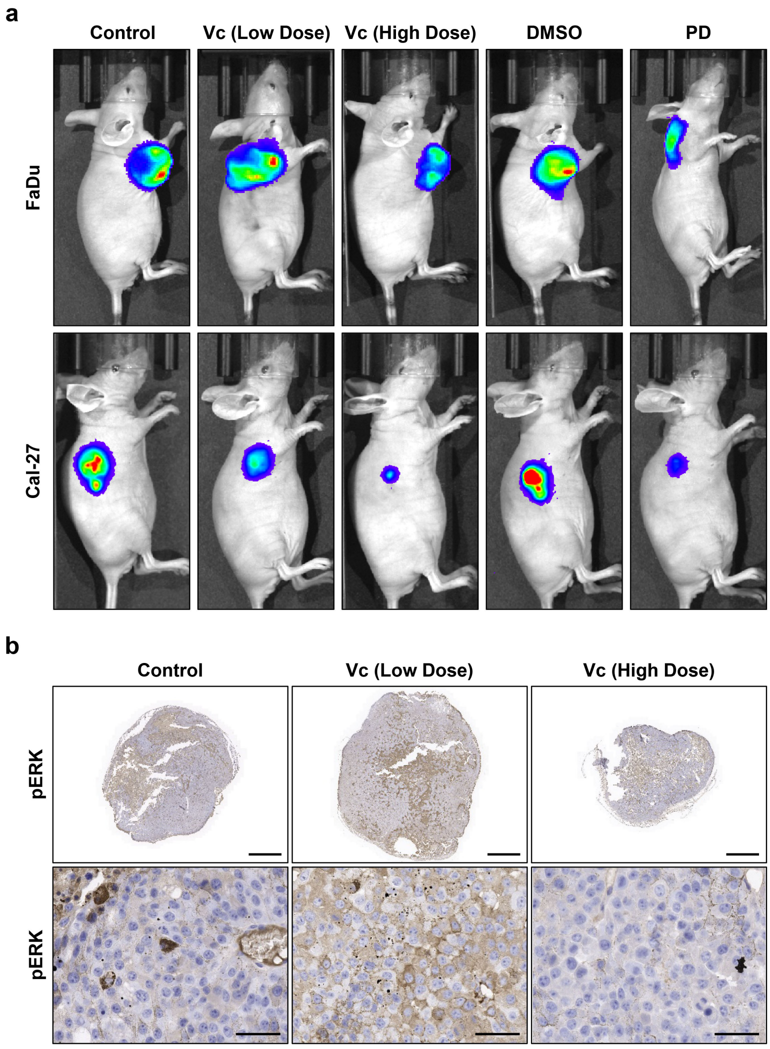


**Figure S9. Vc affects tumorigenesis through ERK** (related to Figure 8)

**(a)** Representative images of in vivo imaging of tumors in mice models constructed by FaDu cells and Cal-27 cells after subcutaneous treatment of Vc or PD.

**(b)** IHC staining of pERK in xenograft tumors constructed by B16-F10 cells. For upper images, scale bar, 3 mm; for lower images, scale bar, 40 μm.

**Table S1. Oligonucleotides used in this study**

| Oligonucleotides | Source | Identifier |
|---|---|---|
| Mus *Svct1* siRNA targeting sequence:<br>GCUCCAUUCUCCUGAUAGUTT | Han et al., 2021 | N/A |
| Mus *Svct2* siRNA targeting sequence:<br>CAGUAUGCCAGAAAUGUUATT | Han et al., 2021 | N/A |
| Human *Svct2* siRNA targeting sequence:<br>GGACUGCGUACAAGUUACAGCUGUU | This paper | N/A |
| The *Ptpn12* sgRNA targeting sequence:<br>forward: 5'-CACCGATATAATCTGAATCTTGGGA-3'<br>reverse: 5'-AAACTCCCAAGATTCAGATTATATC-3' | CCTop-CRISPR/Cas9 target online predictor and CRISPOR | https://cctop.cos.uni-heidelberg.de:8043/ and http://crispor.tefor.net/ |

**Table S2. Antibodies used in this study**

| Antibodies | Source | Identifier |
|---|---|---|
| Ribbit monoclonal anti-pERK1/2 | Cell Signaling Technology | Cat#9101; RRID: AB_331646 |
| Ribbit monoclonal anti-ERK1/2 | Cell Signaling Technology | Cat#4695; RRID: AB_390779 |
| Ribbit monoclonal anti-pJAK2 ($Tyr^{1007/1008}$) | Cell Signaling Technology | Cat#3771; RRID: AB_330403 |
| Mouse monoclonal anti-JAK2 | Santa Cruz Biotechnology | Cat#sc-390539; RRID: AB_2885075 |
| Mouse monoclonal anti-GAPDH | TransGen Biotech | Cat#HC301; RRID: AB_2629434 |
| Mouse monoclonal anti-β-Tubulin | TransGen Biotech | Cat#HC101 |
| Rabbit polyclonal anti-SVCT2 | Abcam | Cat#ab229802 |
| Mouse monoclonal anti-Flag | Sigma-Aldrich | Cat#F1804; RRID: AB_262044 |
| Polyclonal monoclonal anti-HA | ThermoFisher Scientific | Cat#PA1-985; RRID: AB_559366 |
| Mouse monoclonal anti-GFP | Proteintech | Cat#66002-1-Ig; RRID: AB_11182611 |
| Mouse monoclonal anti-phospho-tyrosine | Cell Signaling Technology | Cat#9411; RRID: AB_331228 |
| Mouse monoclonal anti-phospho-serine | Santa Cruz Biotechnology | Cat#sc-81514; RRID: AB_1128624 |
| Mouse monoclonal anti-PTPN12 | Santa Cruz Biotechnology | Cat#sc-271351; RRID: AB_10610603 |
| Goat anti-Mouse IgG (H+L) Highly Cross-Adsorbed Secondary Antibody, Alexa Fluor 555 | ThermoFisher Scientific | Cat#A-21424; RRID: AB_141780 |
| Goat anti-Mouse IgG (H+L) Highly Cross-Adsorbed Secondary Antibody, Alexa Fluor 488 | ThermoFisher Scientific | Cat#A-11029; RRID: AB_138404 |
| Ribbit monoclonal anti-Ki-67 | MXB Biotechnologies | Cat#RMA-0731 |
| Rabbit polyclonal anti-E-Cadherin | Proteintech | Cat#20874-1-AP; RRID: AB_10697811 |
| Ribbit monoclonal anti-N-Cadherin | Cell Signaling Technology | Cat#13116; RRID: AB_2687616 |
| Rabbit polyclonal anti-Vimentin | Proteintech | Cat#10366-1-AP; RRID: AB_2273020 |
| Normal Mouse IgG | Beyotime Biotechnology | Cat#A7028 |
| HRP-labeled Goat Anti-Rabbit IgG (H+L) | Beyotime Biotechnology | Cat#A0208; RRID: AB_2892644 |
| HRP-labeled Goat Anti-Mouse IgG (H+L) | Beyotime Biotechnology | Cat#A0216; RRID: AB_2860575 |

**Table S3. Software and algorithms used in this study**

| Software and algorithms | Source | Identifier |
|---|---|---|
| ImageJ V1.5 | Schneider et al., 2012 | RRID: SCR_003070<br>URL: https://imagej.net/ |
| cellSens Dimension V3.1 | Olympus cellSens Software | RRID: SCR_014551<br>https://www.olympus-lifescience.com/en/software/cellsens/ |
| StepOne software V2.3 | ThermoFisher Scientific | RRID: SCR_014281;<br>https://www.thermofisher.com/us/en/home/technical-resources/software-downloads/StepOne-and-StepOnePlus-Real-Time-PCR-System.html |
| Image Lab V6.0.0 | Bio-Rad Laboratories, Inc. | https://www.bio-rad.com/en-cn/category/image-lab-software-resources?ID=PJWA0VTU86LJ |
| FlowJo V10.0.7 | BD Biosciences | RRID: SCR_008520;<br>https://www.flowjo.com/ |
| GraphPad Prism V6.07 | N/A | RRID: SCR_002798;<br>https://www.graphpad.com/scientific-software/prism/ |
| CaseViewer V2.2 | 3DHistech | N/A |
| FRET analysis (Xia) | Xia and Liu, 2001 | N/A |
